# Magnitude and Kinetics of a set of Neuroanatomic Volume and Thickness together with White Matter Hyperintensity is definitive of Cognitive Status and Brain Age

**DOI:** 10.1101/2023.09.05.556337

**Authors:** Neha Yadav, Niraj Kumar Gupta, Darshit Thakar, Vivek Tiwari

**Affiliations:** Indian Institute of Science Education and Research (IISER) Berhampur, India

## Abstract

In an aging population, a subset of individuals at a given age group have low white matter hyperintensity (WMH) while another subset has intermediate to high WMH load. WMH load quantification together with comprehensive neuroanatomic volumetry needs to be examined together for establishing a unique precise and optimal number of brain features as a noninvasive indicator of ‘Brain Age’ and cognitive status. Here, a comprehensive neuroanatomic volumetry together with WMH quantification using longitudinal MRI and cognitive measurements from two aging cohorts have been performed together with machine learning modeling of the quantitative changes with aging to establish Optimal unique brain events discriminative of cognitive status and estimative of Brain Age. A set of Three optimal brain-associated quantities; wherein two are neuroanatomic features Total brain volume; CSF volume, and the third is the extent of microvascular pathology WMH load, provide highly precise discrimination of cognitive status as cognitively normal (CN), impaired (CI) and AD (CI-AD). While medial cortical thinning of Parahippocampal gyrus is an ‘early age event’ discriminative between CI and CI-AD but loss of hippocampus, gray matter and white matter volume lacks sensitivity to discriminate between CI and CI-AD. The Brain Age estimation using the neuroanatomic volumetry and periventricular and deep WMH load indicates that elevated WMH load in the brain led to an increased Brain Age gap than the brain with low WMH at a given chronological age. Increased Brain Age gap with elevated WMH load at the early age groups is suggestive of profound vascular insult arising from WMH to the brain structure and function.

## Introduction

Aging is one of the major risks for brain disorders^1^. Order, magnitude and kinetics of neuroanatomic and microstructural changes associated with chronological aging of cognitively normal subjects needs to be precisely delineated from that of the events unique to cognitive impairment in MCI and Alzheimer’s disease (AD). Atrophy of various brain structures with chronological aging is not uniform^2–9^. While there is structural atrophy with chronological aging, there is a common phenomenon of demyelination and fiber degeneration due to micro-vessel infarcts, observed as white matter hyperintensity on brain MRI. Given the subtle structural, microstructural and microvascular changes a large fraction of these changes overlaps between normal and aging associated cognitive pathologies. However, the impaired brain health and cognition in MCI and AD may arise upon unique alterations in the magnitude, temporal sequence and kinetics of specific structural and cerebrovascular abnormalities distinct from that observed during chronological aging.

Aging and aging associated cognitive disorders are multifactorial events, therefore, to date, there are no definitive *in vivo* non-invasive quantitative signatures for brain health and determinants of cognitive status as CN, MCI or Alzheimer’s disease^10–16^. A precise measure of the rate of structural atrophy and hypertrophies associated with a set of brain regions together with estimates of white matter hyperintensity with chronological aging would potentially represent overall brain health. In brain MRI investigations of aging subjects, WMH is commonly observed as microvascular pathology, wherein a set of individuals at a given age group have low WMH while another subset has high WMH load^17,18^. Therefore, while aging studies focus on structural alterations, WMH load must be investigated and a comprehensive investigation of brain health must encompass inclusion of WMH and neuroanatomic volumetry.

Here, we have performed comprehensive quantification of brain structures and WMH load, their sequence and temporal order, kinetics and magnitude of alterations with chronological aging, to establish Optimal number of set of brain features and its threshold depictive of Brain Age and discriminative of cognitive status as normal (CN), cognitively impaired (CI), and cognitively impaired with Alzheimer’s disease (CI-AD) using the cohort from National Alzheimer’s Coordinating Center (NACC). Further, we have developed a unique Brain Age (BA) estimation model using the MRI quantified neuroanatomic features together with white matter hyperintensity volume.

## Methods

### Study population and Characteristics

Longitudinal MR Images, clinical investigations, and cognitive status of subjects labeled as cognitively normal (CN), cognitively impaired (CI), and cognitively impaired with an etiological diagnosis of Alzheimer’s disease (CI-AD) were obtained from National Alzheimer’s Coordinating Center (NACC) for the subjects enrolled during 2005 to 2021. NACC is a longitudinal multicenter study established in 1999 by the National Institute of Aging (NIA) which collects and standardizes clinical and neuropathological data from Alzheimer’s Disease Research Centers (ADRCs) across the United States^19–21^. The cognitive status labeled as CN, CI, and CI-AD are based on the NIAA-NIND criterion employed by NACC as a variable NACCALZD. A label of NACCALZD = 8 signifies Cognitively Normal (CN) subjects, NACCLAZD = 0 denotes subjects with cognitive impairment (CI) because of MCI and other disorders, while NACCLAZD = 1 denotes subjects who have cognitive impairment with an etiologic diagnosis of Alzheimer’s disease. Longitudinal T1-weighted (T1w) and T2-FLAIR MRI scans (N = 3058) obtained from the subjects (N = 2114) enrolled in this study till September 2021 from 16 ADRCs (https://www.alz.washington.edu/) were investigated and measured.

Subjects with MRI examination within the range of 1-year from the cognitive measurements were included for brain morphometric measurements. The total number of longitudinal MRI scans investigated in this study across CN, CI, and CI-AD subjects were 2642, wherein, the number of the subjects identified as CN (N_CN_ = 1082), CI (N_CI_ = 197), and CI-AD (N_CI-AD_ = 588) had a total number of 1616, 252, and 774 MRI scans, respectively (Supplementary Table S1).

### Brain region Volumetry and Thickness Quantification

The MRI-based brain region volumetry and thickness were obtained using Imaging of Dementia & Aging (IDeA) Lab pipeline for investigating the chronological aging associated changes in the CN, CI and CI-AD subjects. The brain volumetry quantification, kinetics with aging, and optimization of brain features predictive of cognitive status was trained and tested using machine learning. The anatomical volumes and the rate of change with age for gray matter (GM), white matter (WM), total brain volume (BRNV), hippocampus (HP), lateral ventricles (LV), cerebrospinal fluid (CSF), white matter hyperintensity (WMH) and medial temporal cortical thickness (parahippocampal gyrus: PHG, entorhinal cortex: EC) with aging were investigated across CN, CI and CI-AD subjects. MRI determined volume and thickness across CN, CI, and CI-AD groups was stratified into three age ranges i.e., 50-64 (early), 65-79 (intermediate), and ≥80 (late) as described in earlier investigations^22,23^. The brain MRI regional volume and thickness across the three age groups were used to identify the early, intermediate, and late brain events distinctive of CN, CI, and CI-AD. Number of the data points across the cognitive groups at the early age group are (CN = 412, C I= 33, and CI-AD = 80), at the intermediate age group (CN = 866, CI = 138, CI-AD = 388) and at the late age group (CN = 338, CI = 81, CI-AD = 306).

The regional rate of change of volumes or thicknesses of brain regions with chronological age was determined by a multivariate linear regression^24^ model by setting the intercept at 50 years of age using an R package (R 4.1.2, emmeans 1.8.3). Volume of brain regions obtained from the brain segmentation were normalized with total intracranial volume (ICV) using equation (1):

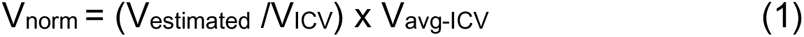

V_norm_ denotes normalised volume; V_estimated_ is the volume obtained from segmentation and V_avg-ICV_ represents mean ICV^25,26^.

### Brain region volume and thickness association with Chronological Aging

A multivariate linear regression model^24,27^ was employed to examine the relationship between the change in brain region volume or thickness with chronological age in CI and CI-AD relative to the CN group, using following equation:

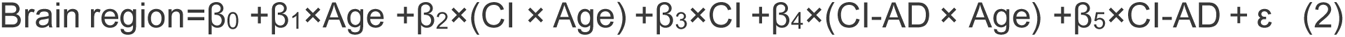

In the equation, the brain region volume or thickness are the dependent variables modeled as a function of several independent variables: Age, and cognitive status (CI, and CI-AD). The coefficients β_0_, β_1_, β_2_, β_3_, β_4_ and β_5_ are the parameters of the model that represent the effect of each independent variable on the dependent variable (Supplementary Table S4). The intercept β_0_ represents the value of the dependent variable when all independent variables are equal to zero. The slope β1 represents the change in the dependent variable per unit change in Age for the control group (CN). The interaction coefficients β_2_ and β_4_ represent the effects of CI (when CI=1) and CI-AD (when CI-AD=1) in addition to the effects of chronological aging, respectively. The β_3_ and β_5_ coefficients represent the effect of CI (when CI=1) and CI-AD (when CI-AD=1), respectively on the brain regions when the age is held constant. The error term ɛ represents additional variabilities in the dependent variable which may not be explained by the effect of the independent variables.

### Kinetics of White Matter Hyperintensity with age

Segmentation of T2w-FLAIR and T1w images using the IDeA lab pipeline provided the total WMH volume. The total WMH volume estimated from the segmentation was normalized by the ICV. An exponential growth curve model using an in-house function was fitted and optimized for investigating the kinetics of WMH progression^28^ with chronological aging across the three cognitive groups, CN, CI and CI-AD as following:

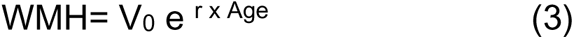

Where, V_0_ is the initial total WMH volume at 50 years of age and r is the rate constant. Additionally, the periventricular WMH (PVWMH) and Deep WMH (DWMH) volume was extracted and calculated from T1-weighted and T2-FLAIR images using a cluster-based pipeline called “UBO Detector”^29^ was used along with brain region volumetry for ‘Brain Age’ estimation.

### Determining Optimal Number of MRI-obtained Brain Features and Cognitive Discrimination Accuracy

A combination of regional brain anatomical volume, cortical thickness, total WMH load obtained from brain MRI segmentation (from IDeA Lab), together with age and gender were used as inputs for machine learning (ML) based supervised platform to obtain a minimum number of ‘Optimal Brain Features’ discriminative of cognitive status as CN, CI, and CI-AD. The scikit-learn library^30^ in Python was employed for executing ML algorithms^31^. Supervised Random Forest method, Simple Classification tree, Bagging Classification, and XGB Classifier were employed to optimize the number of MRI features best suited for identifying CN, CI, and CI-AD groups. A subset of the MRI outputs of regional brain anatomical volume, cortical thickness, total WMH load from CN, CI and CI-AD data was randomly distributed into training and test sets. The training sets were used to train the ML model, while the test set was reserved for evaluating the model’s performance. Due to the inherent imbalance in the number of samples in each cognitive group, a stratified sampling approach was used to ensure that both the training and test sets were representative of the overall dataset, with a similar distribution of class labels and other relevant characteristics.

To assess the accuracy of each ML model and validate their performance for all combinations of MRI features, a stratified k-fold cross-validation technique with k=5 was performed. The cross-validation process was iterated for 6 times to ensure comprehensive and robust evaluation of the models. During each iteration, the algorithm was trained on a random subset of the data. The performance metrics such as accuracy, precision, recall score and confusion matrix analyses were performed to evaluate the performance of the ML models. Also, the accuracy and average normalized confusion matrix resulting from the stratified k-fold cross-validation with 6 repeats were used to assess the stability and the reliability of the cognitive status prediction.

### Determining the Brain Age (BA) from Neuroanatomical structures and WMH

To establish an accurate Brain Age model, a comprehensive segmentation of large number of brain structures and Periventricular white matter hyperintensity (PVWMH) and Deep white matter hyperintensity (DWMH) was carried out in cognitively normal subjects (727 longitudinal MRI scans from 528 subjects) using automated cortical reconstruction segmentation method (FreeSurfer Version 7.2.0)^32^ and UBO Detector pipeline^29^, respectively. The average of the regional brain volume, thickness, and area from the left and right regional values obtained from the FreeSurfer Segmentation were used in the BA model.

Therefore, a total of 178 neuroanatomical features obtained from FreeSurfer segmentation, and periventricular WMH (PVWMH) and Deep WMH (DWMH) volume obtained from the UBO detector together with chronological age (CA) as independent variable were used for establishing a machine learning based model depictive of Brain Age (BA).

The training set involved neuroanatomic volume, thickness, and area from the 221 scans (out of 727 scans) which had undetectable or low WMH (PVWMH <1.5 ml and DWMH <1.5 ml) load from the Cognitively Normal subjects. The nil or low WMH subjects were included to represent any structural and microvascular abnormality free healthy brain subject: a data that rules out any plausible effect of WMH on brain health. Bagging and error correction techniques were performed by dividing the training data into numerous train and validation sets, such that each sample serves as a training set in one of the trials and validation set in another trial. The training and validation processes were iterated for 50 times.

Subsequently, the trained Brain Age model was used to predict Brain Age and Brain Age Gap^33^ (BAG) using the following equation.

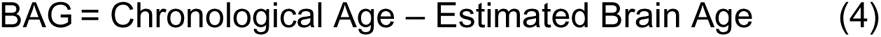

The top ten important features of the brain age prediction model were derived using permutation importance from scikit-learn^30^ python library.

The Brain Age predictive model was also cross validated for cognitively normal subjects, from the ADNI3 cohort (N=92), with Low (0 - 1.5 ml) and High (5 - 10 ml) WMH.

### Statistical Analysis

The dataset was stratified into three age groups (50-64, 65-79, and ≥80) and for each age group the global significance was tested using the nonparametric multivariate model from **npmv** R package^34^. The response variables in the model were brain region volumes and thickness while the factors were the three cognitive status. The **npmv** package computes the F-approximation and permutation p-values for four types rank-based tests: ANOVA type, Wilks’ Lambda type, Lawley Hotelling type, and Bartlett Nanda Pillai type. Additionally, the nonparametric relative effect of cognitive status on each brain region volume and thickness was computed. Furthermore, the subset algorithm was performed based on factor levels (cognitive status) and response variables (brain region volume and thickness) and ANOVA type statistics was chosen as the test statistic. All appropriate subsets, for the three factor levels and eight response variables were checked using multiple testing controlling for familywise error rate at **α** = 0.05. The total number of tests performed for three factor level combinations were 4 (number of factor levels a=3, 2^a^- a - 1= 4) and for sets of eight response variables were 255 (number of variables p = 8, 2^p^ −1 = 255).

In addition, to evaluate differences in brain region volume or thickness between cognitive groups (CN vs CI, CN vs CI-AD, and CI vs CI-AD) for each age group, unpaired two-tailed Welch’s *t*-test was conducted and t-values were obtained. Bonferroni correction was applied to account for multiple comparisons, thereby the significance threshold for *t*-values were set at p<0.016.

The multivariate linear regression (equation (2)) for brain region volume or thickness was performed using the emmeans library in R (R 4.1.2, emmeans 1.8.3) and the significance of the full model was tested using the *f*-test. The ‘p’ values <0.05 were considered significant for the coefficients β_0_, β_1_, β_2_, β_3_, β_4_, and β_5_ obtained from equation (2).

The Supervised Machine Learning models were trained using various combinations of MRI-derived neuroanatomic structures to discriminate the cognitive status. The trained machine learning models were evaluated for each combination of brain features using accuracy, precision and recall analysis to assess the performance of the model in correctly classifying subjects as CN, CI, and CI-AD.

## Results

### Brain Structure Morphometry Measurements with Aging Across Cognitive Groups

The nonparametric global multivariate analysis (ANOVA type) examining the neuroanatomic differences between CN, CI and CI-AD revealed significant differences for gray matter, white matter, hippocampus, lateral ventricle, CSF, total brain volume and medial temporal lobe cortical thickness (entorhinal cortex thickness, and parahippocampal gyrus thickness) at the early age (F = 22.31, p = 0), intermediate age (F = 41.39, p = 0) and late age (ANOVA type test, F = 16.33, p =0) (Supplementary Table S2A, B). At the early age group (50-64) all the eight brain volumetry quantities showed a significant global difference across the cognitive groups while at the intermediate and late age group, no global difference was obtained for the white matter volume. Furthermore, unpaired two-tailed Welch’s *t*-test revealed the extent and pattern of differences between two cognitive groups for a given brain region at early, intermediate and late age groups. In addition to the structural volumetry, tests were performed for ascertaining age stratified differences for white matter hyperintensity load across cognitive groups. A combined approach of structural volumetry and white matter hyperintensity load were used together for the machine learning based investigations for cognitive status discrimination and brain age estimation.

### Gray Matter, White Matter and Total Brain Volume with Aging in CN, CI vs CI-AD

Whole brain segmentation of T1-weighted images revealed a progressive loss in gray matter, white matter and total brain volume with age in cognitively normal (CN), cognitively impaired (CI), and cognitively impaired with Alzheimer’s disease (CI-AD) subjects. At the early age group, the gray matter, white matter and total brain volume was significantly lower in CI and CI-AD subjects compared to the CN (Fig.1 A, C, Fig.S1 A, B, Fig.S2 A). The t-map depicted by the t-values suggests progressive loss in GM and WM with early atrophy in CI and CI-AD subjects marked with progressive loss with aging (Fig.1 A, Fig.S1 A). Significant loss in the gray matter volume in the CI (−5.2%) and CI-AD (−5.6%) subjects was observed at the early age compared to the CN (Fig.1 C, Supplementary Table S3). However, there was no significant difference in the gray matter volume between CI and CI-AD for all the three age groups (Fig.1 C, Supplementary Table S3, S5). The multivariate linear regression model as described in equation-2 (Supplementary Table S4) revealed loss of −2.1 ml of gray matter volume for CN subjects per year, while CI and CI-AD subjects showed lower slope of decline (CI: −1.4 ml, p_(CN_ *_vs_* _CI)_ = 0.0002; CI-AD: −1.5 ml, p_(CN_ *_vs_* _CI-AD)_ = 2.2×10^-6^), by a factor of −33% and −28%, respectively compared to the CN (Fig.1 E). Whereas, the baseline volume of gray matter in CI and CI-AD subjects was remarkably lower compared to the CN group as observed from the intercepts (obtained upon restricting the initial age to 50 years) in the multivariate linear regression (GM-I_(CN)_ = 636.2 ml, I_(CI)_ = 608.2 ml, I_(CI-AD)_ = 606.9 ml, Intercept-GM: p_(CN_ *_vs_* _CI)_= 1×10^-7^, p_(CN_ *_vs_* _CI-AD)_ < 2×10^-16^, p_(CI_ *_vs_* _CI-AD)_= 0.83).

**Figure 1.**
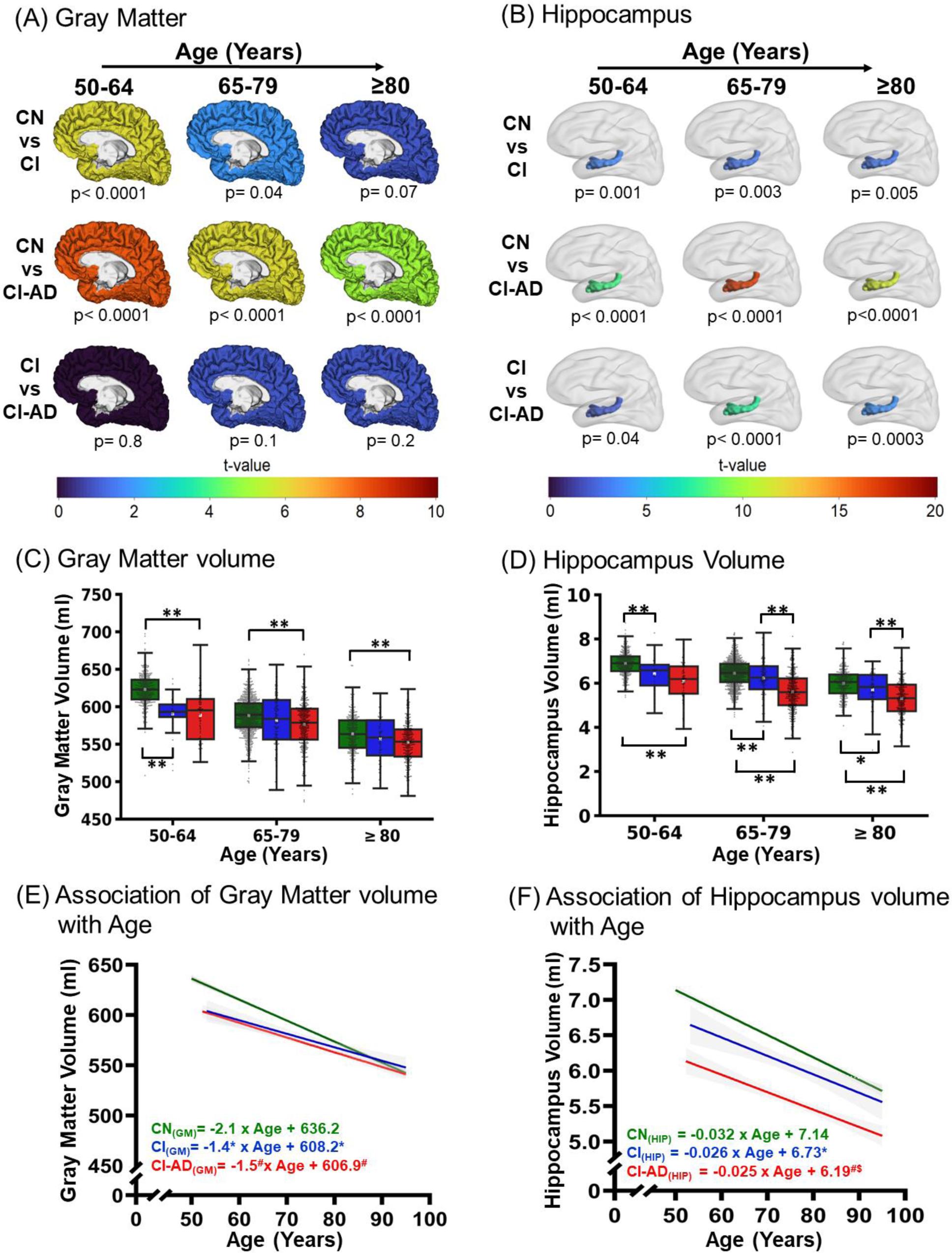
Magnitude and kinetics of Gray matter and Hippocampus volume with age across CN, CI, and CI-AD subjects. **(A)** Gray matter and **(B)** Hippocampus volume comparison between CN *vs* CI, CN *vs* CI-AD, and CI *vs* CI-AD subjects depicted by the t-map across three age groups 50-64 (early), 65-70 (intermediate), and ≥80 (late), wherein warmer color depicts remarkable difference in the gray matter and hippocampus volume between the two cognitive groups at the given age range. The significance was set to p <0.016 upon Bonferroni correction. The color bar depicts the *t-*value. Boxplot showing the median and mean **(C)** gray matter and **(D)** hippocampus volume in CN, CI, and CI-AD subjects. The gray matter and hippocampus volume between the cognitive groups was compared using unpaired, two-tailed Welch’s *t*-test followed by Bonferroni correction. The upper margin of the box plot represents the Q3 (third quartile), and the lower margin of the box represents the Q1(first quartile). The height of the box represents the interquartile range (IQR); the median is represented by the black line inside the box and the white square in the middle represents the mean of the sample. Statistical significance for comparing the mean gray matter volume between the cognitive groups (CN, CI, and CI-AD) across the stratified age groups is depicted as * p <0.016, **p <0.001. Linear regression of volume of **(E)** gray matter (GM) and **(F)** hippocampus volume with age. The linear regression was performed upon setting up the age intercept at 50 years of age for all the three cognitive groups. Green, Blue, and Red represent CN, CI, and CI-AD respectively. Statistical significance for the slope and intercept comparison between CN vs CI (*), CN vs CI-AD (^#^), and CI vs CI-AD (^$^) was set at p< 0.05.

Similarly, progressive loss in white matter was observed with age for all the three cognitive groups. Significantly reduced mean white matter volume was noted at the early age group for both CI and CI-AD compared to the CN subjects (CN: 493.2 ml, CI: 474.6 ml, CI-AD: 474.8 ml). The differences between the groups get masked at intermediate and late age groups (Fig.S1 B). Moreover, CI and CI-AD subjects had similar white matter volume loss compared to CN at the early age group. The annual white matter loss with age for the CN group was −3.3 ml, while a significantly slower rate of annual decline in the white matter - 26% and −24.8% (p_(CN_ *_vs_* _CI)_ = 0.0008, p_(CN_ *_vs_* _CI-AD)_ = 9.2×10^-7^, p_(CI_ *_vs_* _CI-AD)_ = 0.91) was observed for the CI and CI-AD groups, respectively (Fig.S1 C). The multivariate regression showed that the baseline volume of white matter for CI and CI-AD group was remarkably lower compared to the CN group (WM-I_(CN)_ = 516.6 ml, I_(CI)_ = 500.4 ml, I_(CI-AD)_ = 498.0 ml, Intercept-WM: p_(CN vs CI)_ =0.015, p_(CN vs CI-AD)_ = 3.3×10^-5^, p_(CI vs CI-AD)_ = 0.74).

Similarly, the per year loss of the total brain volume (= gray matter volume + white matter) was −5.4 ml for the CN group, while a significantly slower kinetics was observed for the CI (−30%) and CI-AD (−27%) groups compared to CN group (p_(CN_ *_vs_* _CI)_ = 1.1×10^-8^, p_(CN_ *_vs_* _CI-AD)_ = 2×10^-16^, p_(CI_ *_vs_* _CI-AD)_ = 0.61) (Fig.S2 B). Noticeably, the baseline volume of total brain was significantly lower for the CI and CI-AD groups compared to the CN (I_(CN)_ = 1152.9 ml, I_(CI)_ = 1108.6 ml, I_(CI-AD)_ = 1104.9 ml, Intercept: p_(CN_ *_vs_* _CI)_ = 1.5×10^-9^, p_(CN_ *_vs_* _CI-AD)_ < 2×10^-16^, p_(CI_ *_vs_* _CI-AD)_ =0.65) (Fig.S2 B).

### Hippocampus volume kinetics with Aging in CN CI, and CI-AD

A significant reduction in the hippocampus volume was observed with age across CN, CI, and CI-AD subjects. The mean hippocampus volume was significantly lower in the CI and CI-AD subjects compared to the CN group across all the age groups (Fig.1 B, D, Supplementary Table S3, S5). Although, hippocampus volume was not discriminative of CI and CI-AD subjects at the early age group, a significantly distinct hippocampal volume was observed at intermediate (CI: 6.2 ml; CI-AD: 5.6 ml, p=2.1×10^-12^) and late age groups (CI: 5.7 ml; CI-AD: 5.3 ml, p=0.0003), wherein the hippocampus volume of CI-AD was significantly lower compared to CI (Fig.1 B, D).

Unlike the rate of loss of GM, WM and total brain volume, the loss in hippocampal volume with age was similar for the CN, CI, and CI-AD groups (Fig.1 F, Supplementary Table S4). The multivariate regression showed that the baseline hippocampus volume was significantly different across the three cognitive groups as depicted by the intercept (I_(CN)_ = 7.1 ml, I_(CI)_ = 6.7 ml, I_(CI-AD)_ = 6.2 ml). The baseline volume of CI and CI-AD subjects was significantly lower compared to the CN subjects (p_(CN_ *_vs_* _CI)_ = 0.004, p_(CN_ *_vs_* _CI-AD)_ <2×10^-16^). Moreover, the baseline volume of CI-AD was significantly lower than CI (p_(CI_ *_vs_* _CI-AD)_ = 0.0006) subjects (Fig.1 F).

### Medial temporal lobe cortical thinning with Aging in CN, CI, and CI-AD

The medial temporal lobe cortices, entorhinal cortex and parahippocampal gyrus exhibited progressive thinning with age. The entorhinal cortical thickness was significantly lowest in the CI-AD subjects at all the age groups compared to CN and at intermediate and late age group compared to CI(Supplementary Table S3) (Fig.2 A, B). With progression in age, the mean thickness of CI was observed to be relatively reduced compared to CN at the intermediate age group (Fig.2 B). Further, the multivariate linear regression (Supplementary Table S4) of the entorhinal cortical thickness with age for the cognitively normal group showed a thinning rate of −0.005 mm per year (p = 0.0018). The annual reduction of entorhinal cortex thickness with age was ∼3-times faster in CI and CI-AD subjects compared to the CN subjects (p_(CN_ *_vs_* _CI)_ =0.036, p_(CN_ *_vs_* _CI-AD)_ =0.0007) despite the baseline thickness of entorhinal cortex was significantly lower for CI-AD subjects by - 12.3% and −12.8% compared to CN (p_(CN_ *_vs_* _CI-AD)_ = 1.6×10^-8^) and CI subjects (p_(CI_ *_vs_* _CI-AD)_ = 0.001) respectively (Fig.2 D). However, the per year decrease in the entorhinal cortex thickness was comparable in CI and CI-AD subjects (p_(CI_ *_vs_* _CI-AD)_ =0.93) (Fig.2 D).

**Figure 2.**
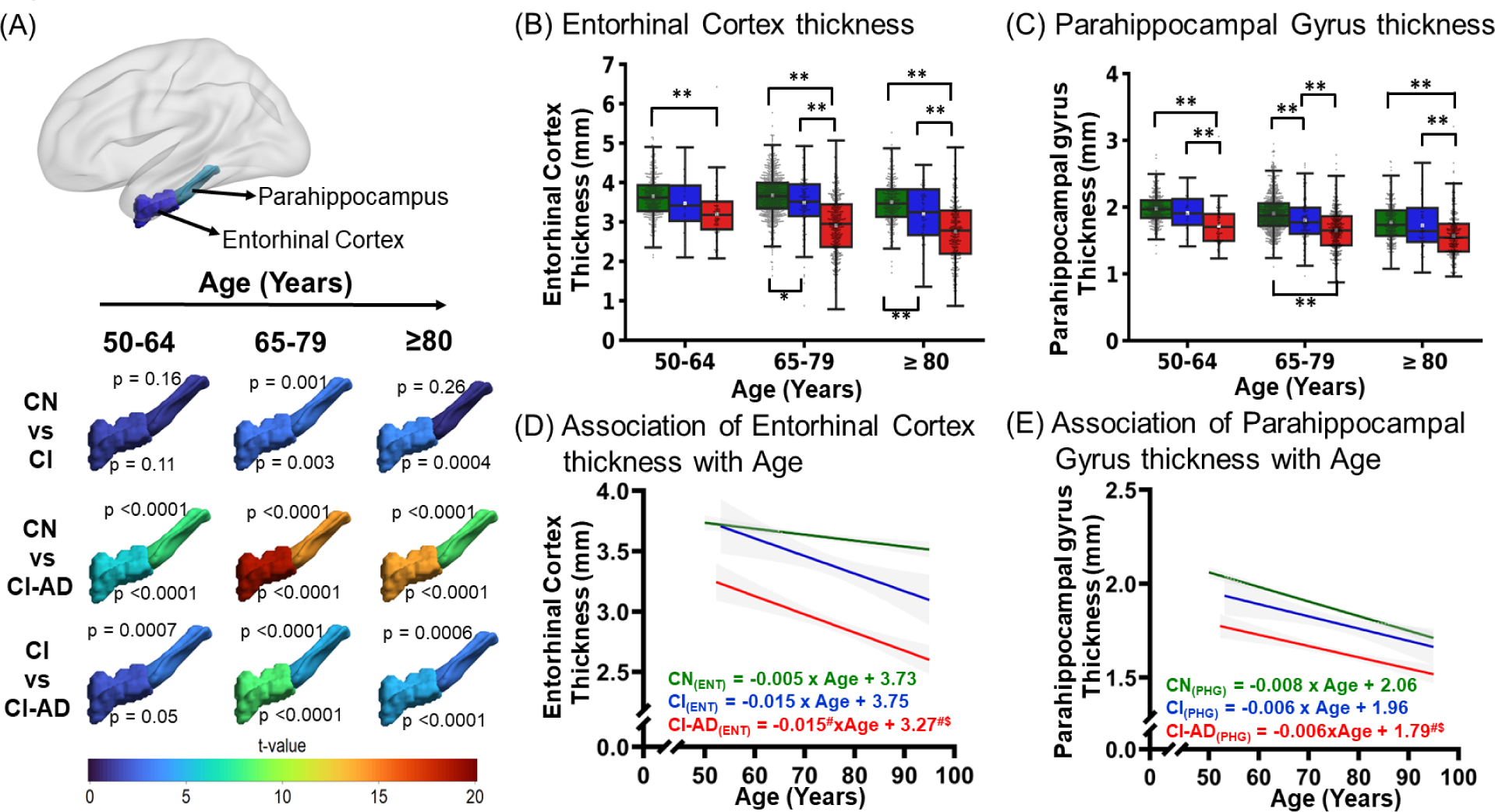
Cortical thinning with age across CN, CI, and CI-AD subjects. **(A)** The t-map of Entorhinal cortex thickness and Parahippocampal gyrus thickness between CN *vs* CI, CN *vs* CI-AD, and CI *vs* CI-AD across early, intermediate, and late age groups. The significance was set to p <0.016 (Bonferroni corrected) and the color bar depicts the *t*-value. Higher the t-value greater the difference between the thickness of cognitive groups. **(C, D)** The Boxplot with median (solid line) and mean (white square) thickness of the entorhinal cortex and parahippocampal gyrus stratified across the early, intermediate, and late age groups for CN (green), CI (blue), and CI-AD (red) subjects. P-values were calculated with the unpaired, two-tailed Welch’s *t*-test followed by Bonferroni correction. Statistical significance for the mean entorhinal cortex and parahippocampal gyrus thickness comparison among cognitive groups (CN, CI, and CI-AD) across the age groups is depicted as * p <0.016, **p <0.001. **(E, F)** Linear regression of entorhinal cortex and parahippocampal gyrus thickness with age upon setting up the intercept at 50 years. Statistical significance for the slope and intercept comparison between CN vs CI (*), CN vs CI-AD (^#^), and CI vs CI-AD (^$^) was set at p <0.05.

Parahippocampal gyrus (PHG) thinning with age revealed a significantly unique aging pattern, wherein CI-AD subjects exhibit reduced thickness compared to CI and CN at early as well as intermediate age groups (Fig.2 A, C, Supplementary Table S3). Additionally, thinning of PHG was discriminative of CI from CN subjects at intermediate age group (Fig.2 C). Although the baseline thickness of parahippocampal gyrus for CI-AD subjects was significantly lower by −13% and −8.6% compared to CN (p_(CN_ *_vs_* _CI-AD)_ = 1.7×10^-13^) and CI subjects (p_(CI_ *_vs_* _CI-AD)_ = 0.008) respectively (Fig.2 E), the yearly rate of PHG thinning with age was similar for all three cognitive groups (CN: 0.008 mm/year; CI:0.006 mm/year; CI-AD: 0.006 mm/year) (Fig.2 E, Supplementary Table S4).

### Lateral Ventricle and CSF volume with Aging in CN, CI, and CI-AD

The lateral ventricle and the CSF volume increased significantly with age for all the three cognitive groups (CN, CI, and CI-AD). Interestingly, the mean lateral ventricle volume of CI-AD group was significantly higher compared to the other two cognitive groups, CN and CI, at the early (CN: 19.3 ml, CI: 27.2 ml, CI-AD: 39.8 ml), intermediate (CN: 29 ml, CI: 38.8 ml, CI-AD: 44.9 ml), and late age groups (CN: 39.5 ml, CI: 44.1 ml, CI-AD: 53.2 ml). Noticeably, CI had significantly higher ventricular volume, distinct from the CN at early as well as other age groups (Fig.3 A, B, Supplementary Table S3).

**Figure 3.**
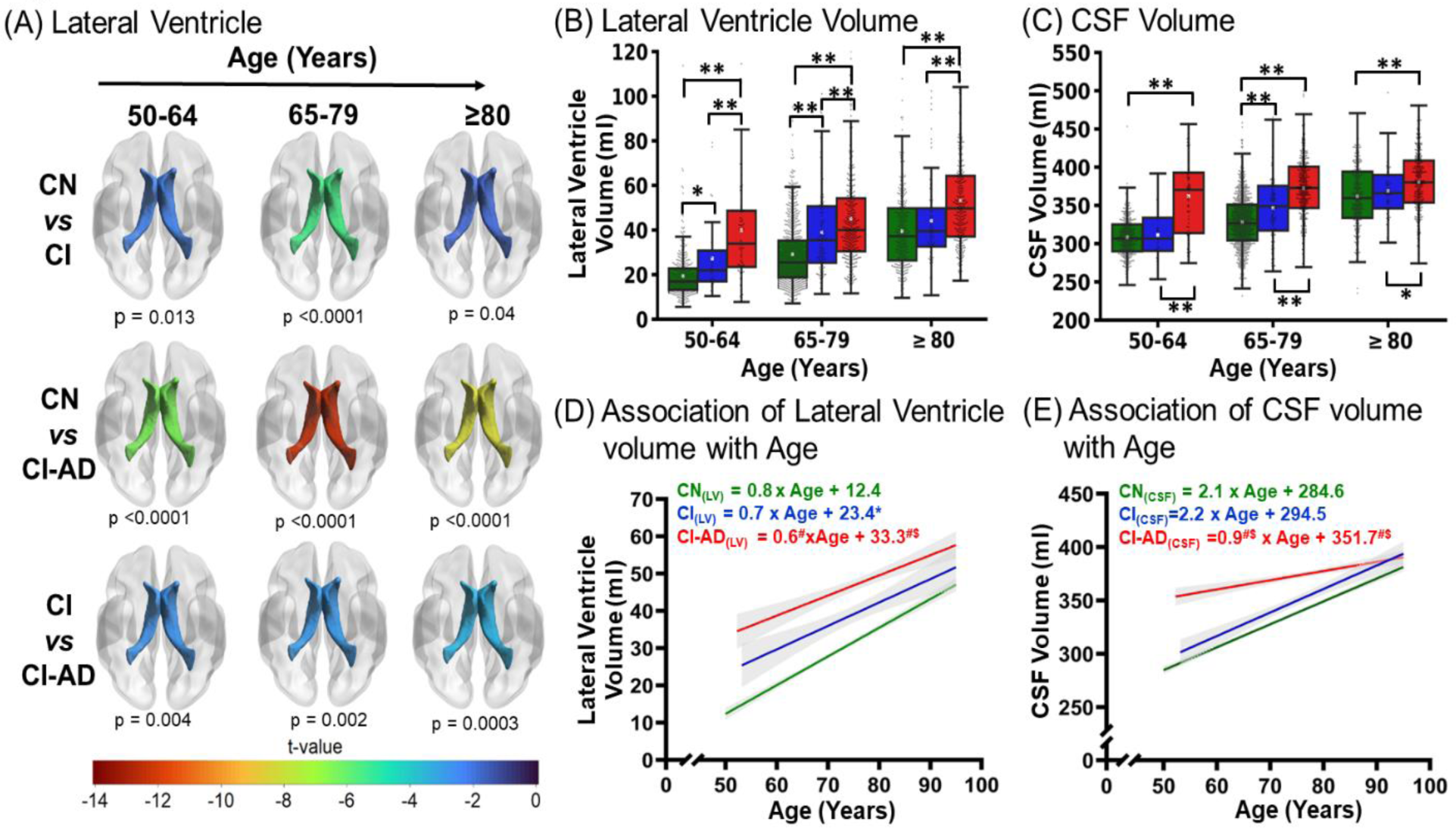
Lateral Ventricular hypertrophy and CSF increase with age across CN, CI, and CI-AD subjects. **(A)** The t-map illustrating the difference of Lateral ventricle volume between CN *vs* CI, CN *vs* CI-AD, and CI *vs* CI-AD across three age groups i.e., 50-64 (early), 65-70 (intermediate), and ≥80 (late). Lower the t-value higher is the difference in the lateral ventricle volume between the cognitive groups. The t-value significance was set at p <0.016 (Bonferroni corrected) and the color bar depicts the *t*-value. Early enlargement of the ventricles is observed for CI and CI-AD groups compared to the CN. Also, early enlargement of ventricles is observed for CI-AD subjects compared to CI subjects. **(B, C)** Lateral Ventricle and CSF volume of CN, CI, and CI-AD subjects across early, intermediate, and late age groups respectively. P-values were calculated with the unpaired, two-tailed Welch’s *t*-test followed by Bonferroni correction. Statistical significance for comparing the mean lateral ventricle and CSF volume among cognitive groups (CN, CI, and CI-AD) across the stratified age groups is depicted as * p <0.016, **p <0.001. **(D, E)** Linear regression of Lateral ventricle volume and CSF shows a progressive increase in the volume across three cognitive groups-CN (green), CI (blue), and CI-AD (red). The linear regression was performed upon setting up the age intercept at 50 years. Statistical significance for the slope and intercept comparison between CN vs CI (*), CN vs CI-AD (^#^), and CI vs CI-AD (^$^) was set at p <0.05.

The annual rate of ventricular increase was 0.8 ml/year for the CN subjects while it was slower by −12% and −25% for CI and CI-AD subjects (p_(CN_ *_vs_* _CI)_ = 0.27, p_(CN_ *_vs_* _CI-AD)_ = 0.0059, p_(CI_ *_vs_* _CI-AD)_ =0.53). The slower ventricular increase in CI and CI-AD subjects, may be potentially attributed to relatively higher intercept for the ventricular volume for the CI and CI-AD subjects compared to the CN (I_(CN)_ = 12.4 ml; I_(CI)_ = 23.4 ml; I_(CI-AD)_ = 33.3 ml, Intercept: p_(CN_ *_vs_* _CI)_ = 0.0009, p_(CN_ *_vs_* _CI-AD)_ < 2×10^-16^, p_(CI_ *_vs_* _CI-AD)_ =0.008) at the baseline age (Fig.3 D, Supplementary Table S4).

The mean CSF volume was significantly higher for CI-AD subjects compared to CN and CI subjects at all the three age groups (Fig.3 C, Supplementary Table S3). Furthermore, the mean CSF volume was significantly distinct between CI and CI-AD (CI: 346.7 ml, CI-AD: 372 ml; p=1.7×10^-9^) as well as between CN and CI (CN: 328.3 ml, CI: 346.7 ml; p=2.3×10^-6^) at the intermediate age group.

The multivariate regression revealed that the per year increase in CSF was significantly slower for the CI-AD group (0.9 ml/year) compared to CN (2.1 ml/year) and CI (2.2 ml/year) groups marked with significantly higher CSF volume in the CI-AD as determined from the intercept (I_(CN)_ = 284.6 ml; I_(CI)_ = 294.5 ml; I_(CI-AD)_ = 351.7 ml) compared to CN and CI groups (Intercept: p_(CN *vs* CI)_ = 0.15, p_(CN *vs* CI-AD)_ < 2 x10^-16^, p_(CI *vs* CI-AD)_ =2.2 x10^-13^) at the baseline age group (Fig.3 E, Supplementary Table S4).

### White Matter Hyperintensity with Aging in CN, CI and CI-AD Groups

T2-FLAIR segmentation showed that the total white matter hyperintensity (WMH) load (deep + periventricular WMH) increases with aging across all three cognitive groups (Fig.4 A). Although the WMH load at the early age group was miniscule and not significantly distinct between CN (1.7 ml), CI (2.3 ml) and CI-AD (2.8 ml) groups, but with progression in age, at the intermediate and late age group higher WMH load (∼2X) was quantified in both CI and CI-AD subjects compared to the CN subjects (Fig.4 B).

**Figure 4.**
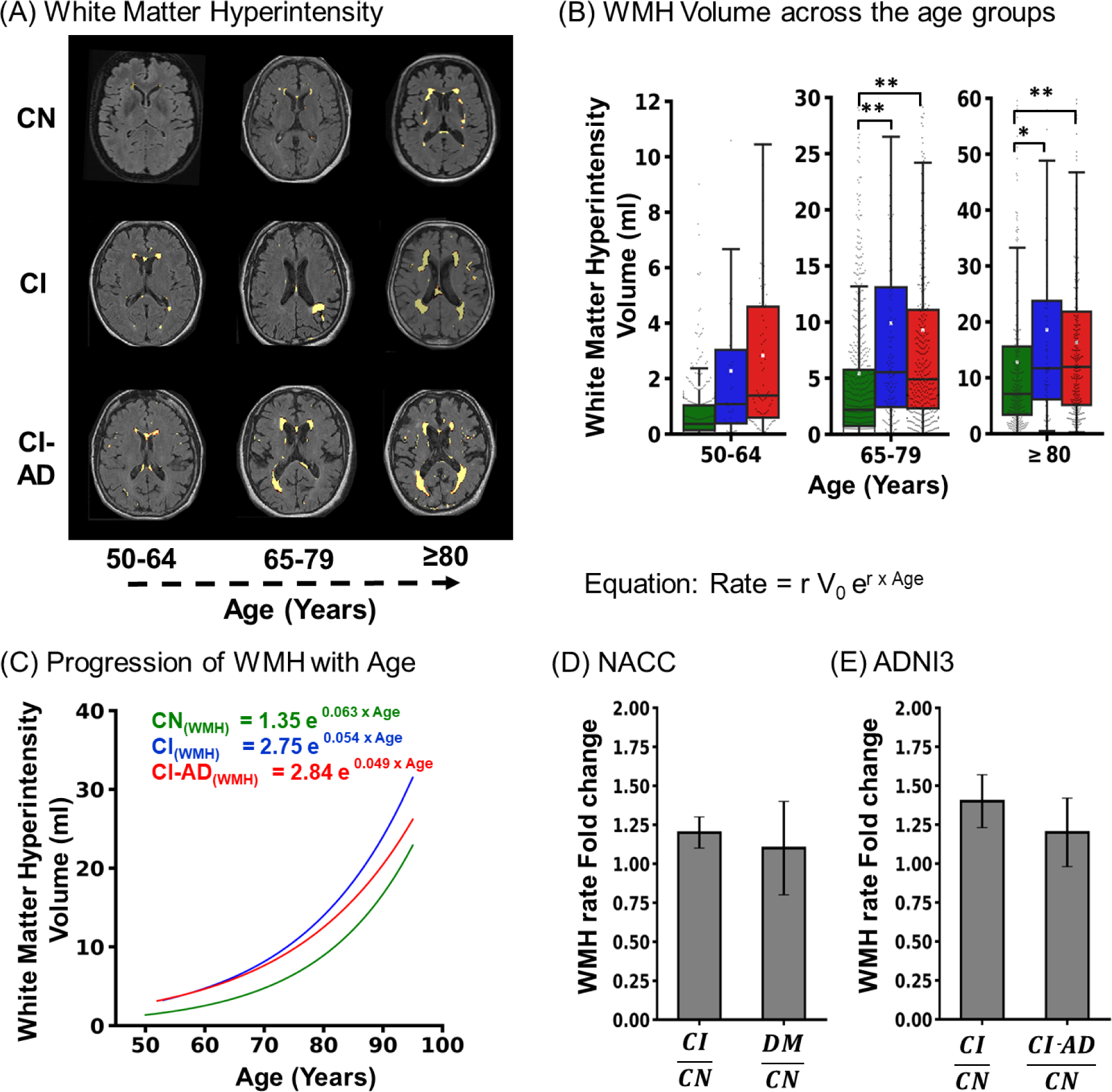
White matter hyperintensity (WMH) loads across CN, CI, and CI-AD subjects. **(A)** Segmentation mask of WMH load was generated from T1w, and T2-FLAIR MRI using Lesion Segmentation Tool (LST) for CN CI and CI-AD subjects across early, intermediate, and late age groups. **(B)** The Boxplot depicts the median (solid line) and the mean (white square) WMH volume across 50-64), 65-79, and ≥80 subjects. P-values were calculated with the unpaired, two-tailed Welch’s *t*-test followed by Bonferroni correction. Statistical significance for the mean WMH comparison among cognitive groups (CN, CI, and CI-AD) across the age groups is depicted as *p <0.016, **p <0.001. **(C)** The exponential increase of total WMH load with age across CN (green), CI (blue), and CI-AD (red) subjects. The equation represents the rate of change of WMH load where, r is the rate constant and V0 is the initial WMH volume at 50 years of age. **(D)** Bar plot depicting mean ± standard deviation WMH rate fold change for CI and CI-AD subjects with respect to cognitively normal (CN) subjects in NACC cohort. **(E)** Bar plot depicting Average WMH rate fold change in CI and Dementia (DM) subjects with respect to CN subjects in ADNI cohort.

The increase in total WMH with age followed the exponential growth pattern wherein the exponential fitting (Fig.4 C) depicted WMH load increase with age for all the three cognitive groups. The WMH initial load observed for CN, CI, and CI-AD was V_0(CN)_ = 1.35 ml, V_0(CI)_ = 2.75 ml, and V_0(CI-AD)_ = 2.84 ml and the rate constant was r_(CN)_ =0.063, r_(CI)_ =0.054, r_(CI-AD)_ =0.049 respectively. WMH load increased in CI and CI-AD subjects across the age groups but there was no significant difference between WMH accumulation and kinetics between the CI and CI-AD groups. At every given age for all the subjects, the rate of increase of WMH load for CI and CI-AD subjects was ∼1.4 times faster (CI/CN: 1.4 ± 0.2; CI-AD/CN: 1.2 ± 0.2) compared to the CN (Fig.4 D).

Similarly, in the ADNI cohort, the rate of increase of WMH load for CI and dementia (DM) subjects was ∼1.2 times faster (CI/CN: 1.2 ± 0.1; DM/CN: 1.1 ± 0.3) compared to the CN (Fig.4 E) similar to that observed in the NACC cohort.

### Machine Learning Method for Optimizing Brain MRI events distinctive of CN, CI and CI-AD subjects

Using the neuroanatomic volumetry and thickness quantifications, WMH volume estimates together with age and gender variables, supervised machine learning algorithms: simple classification tree, random forest, bagging classification, and XGB classifier (Fig.5 A) were trained to obtain the optimal number of brain MRI segmented quantities discriminative of cognitive status.

**Figure 5.**
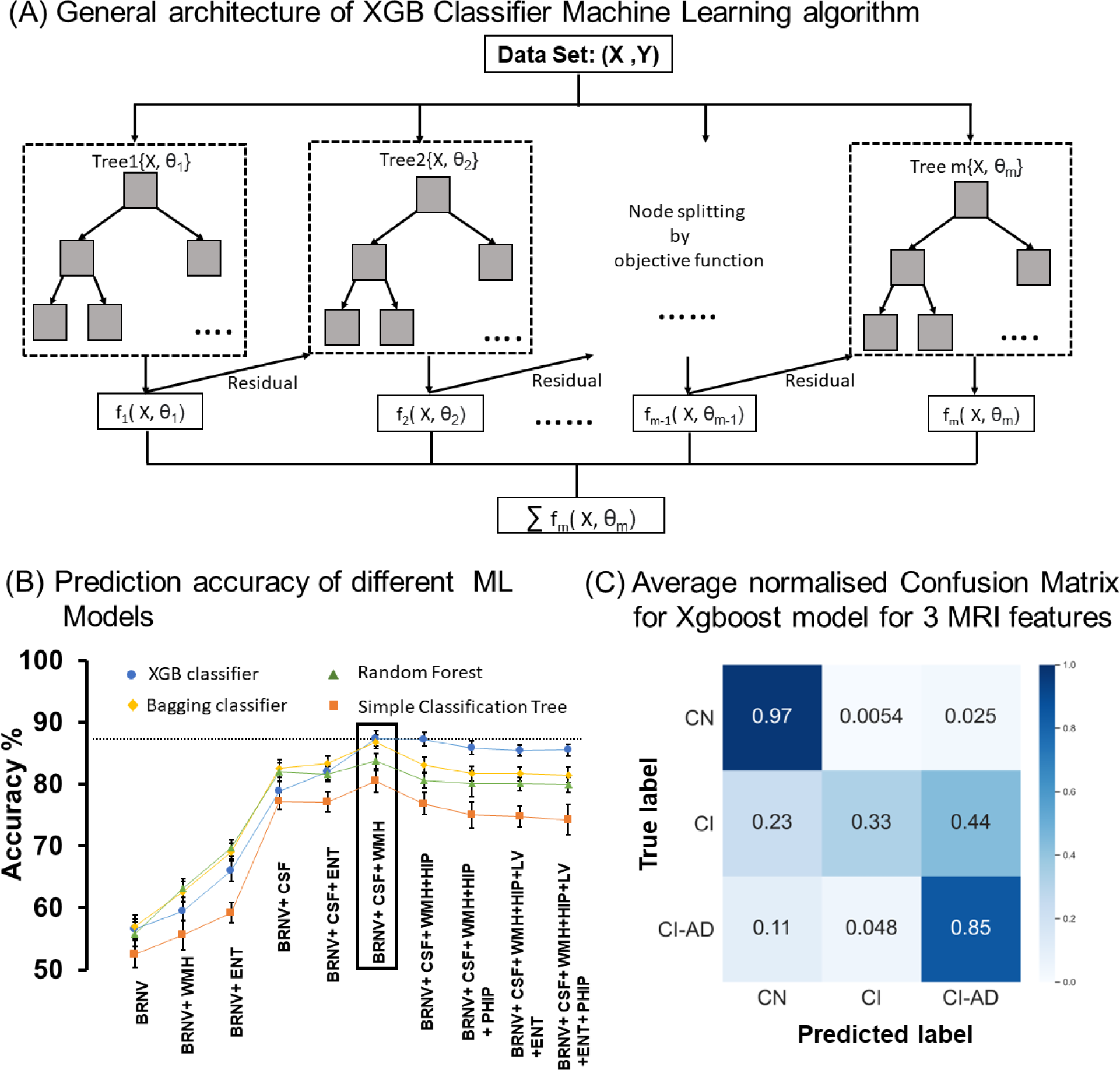
Summary of the Machine learning algorithm performance in predicting the cognitive status of the participants based on the MRI segmented brain volume and thickness and age. **(A)** Schematics of XGB classifier ML model which is based on the gradient boosting technique. **(B)** Cognitive status prediction accuracy of different ML Models for combination of different MRI obtains neuroanatomic volumes and thicknesses. MRI features are added one by one with age and gender to check for the increase in average accuracy of XGB classifier (blue circle), Random Forest (green triangle), Bagging classifier (yellow diamond) and Simple Classification Tree (orange square) ML models. The result shows the mean accuracy ± SD (standard deviation). The highest accuracy for all ML models was obtained for the combination of 3 MRI features *i.e*., total Brain volume, CSF and WMH with age and gender and out of the 4 ML models XGB classifier gave the highest accuracy. **(C)** Average normalized confusion matrix for XGB classifier ML machine models for predicting the cognitive status of the test data for the 3 optimized MRI features (total Brain volume, CSF and WMH) with age and gender which is giving the highest accuracy. **\*** BRNV=Total Brain Volume, CSF=cerebrospinal fluid, LV=Lateral Ventricle, HIP= Hippocampus, WMH= White Matter Hyperintensity, ENT=Entorhinal cortex and PHIP-Parahippocampal Gyrus

Unique predictive accuracy for the Cognitive status as CN, CI and CI-AD was observed upon various combinations of neuroanatomic structures and WMH load. Inclusion of only a single neuroanatomic feature provided accuracies ranging between ∼50-60%, whereas random addition of neuroanatomic volume and WMH discriminates cognitive status with varied accuracies (50 - 88%) (Fig.5 B). A combination of two neuroanatomical features *viz.* Total Brain Volume and CSF together with WMH volume provided the highest average accuracy for cognitive status discrimination using the XGB Classifier (∼87.2 ± 1.3%) (Fig.5 B, C) and Bagging Classification (86.9 ± 1.2 %) (Fig.5 B). Moreover, the combination of these three unique MRI segmented quantities also yields the highest accuracy with other ML algorithms, such as the Simple Classification tree achieved an average accuracy of 80.7 ± 1.9 %, while Random Forest achieved average accuracy of 83.7 ± 1.2 % (Fig.5 B).

### Establishing Brain Age using 180 MRI determined Volumetry Features

Boosting algorithm method was employed to obtain the optimal and accurate architecture model for determining Brain Age from a comprehensive 178 neuroanatomic structures and two microvascular pathology parameters in form of periventricular WMH load and Deep white matter hyperintensity load (DWMH) together with chronological age from the Cognitively normal subjects (Fig.6 A). The training subjects had PVWMH and DWMH as nil or <1.5 ml. Estimates of brain age correlated significantly with chronological age as depicted by the average correlation coefficient (r) as 0.89 ± 0.03 for 50 iterations (Fig.6 B).

**Figure 6.**
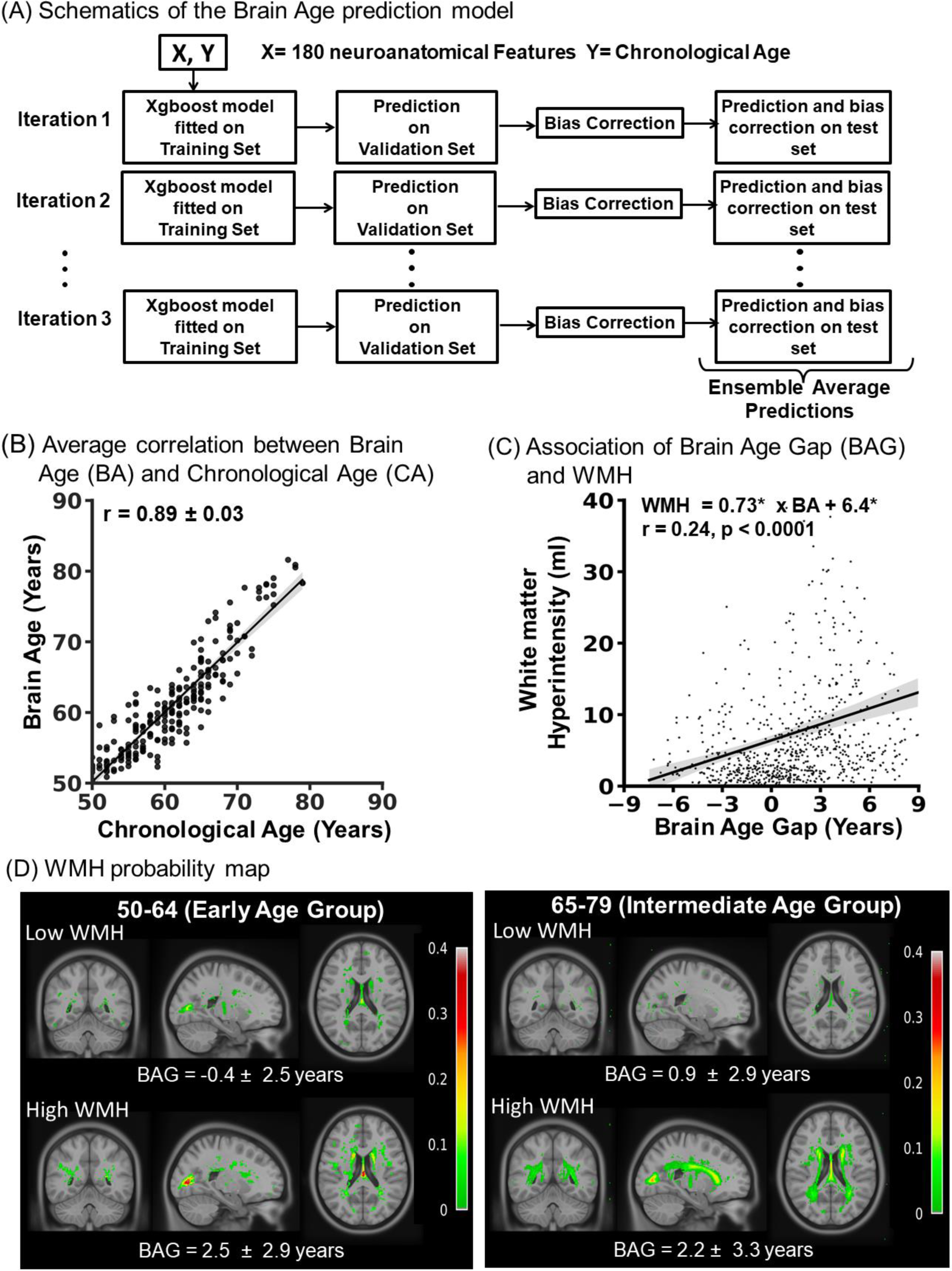
Brain age model developed from 180 MRI obtained neuroanatomical features. **(A)** Schematics illustrating the workflow of the Brain age prediction model. **(B)** The average association between Brain Age (BA) and Chronological Age (CA) obtained from the ensemble average prediction for 50 iterations. **(C)** The linear regressions showing the association of white matter hyperintensity with Brain Age Gap (BAG) for NACC cohort. **(D)** Voxel-wise probability map depicting the occurrence of total WMH load (PVWMH +DWMH) on the coronal, sagittal and axial slice for cognitively normal subjects in the early age group with no or low WMH (WMH <1.5 ml, n=95) and high WMH (WMH 5-10 ml, n=32) and intermediate age group with no or low WMH (n=35) and high WMH (n=118) with mean Brain Age Gap (BAG) and standard deviation.

Examination of increased WMH load on the Brain Age revealed that the Brain Age Gap (BAG) correlates positively with the total WMH load (r = 0.24, p<0.0001) (Fig.6 C). The mean Brain Age Gap for cognitively normal subjects with undetectable or very low WMH <1.5 ml was estimated to be insignificant across the age groups; at the early (−0.4 ± 2.5 years) and intermediate (0.9 ± 2.9 years) age groups. However, for the cognitively normal subjects who had high WMH (5-10 ml) load in the brain, a significantly higher BAG was estimated at the early (2.4 ± 2.9 years) and intermediate (2.2 ± 3.3 years) age groups with (Fig.6 D, Fig.S3 A). Cross validation for the subjects from the ADNI-3 cohort revealed similar BAG estimates as for the NACC cohort for the no and high WMH subjects (Fig.S3 B).

Permutation importance analyses for the top ten brain volumetry contributors towards Brain age estimation for the subjects with no WMH, revealed that volume of 3rd ventricle is the topmost feature with an importance factor of 0.14 (Fig.S4 A) while, for the subjects with high WMH, the volume of periventricular white matter hyperintensity (PVWMH) was the most important feature with an importance factor of 0.083 (Fig.S4 B). However, a set of brain structures; volume of accumbens area, ratio of brain segmented volume by total intracranial volume, and amygdala are common top features contributing towards brain age for both the groups *viz* no WMH and high WMH, but the extent of contribution is distinct. It is noticeable that in the case of high WMH subjects, PVWMH contributes maximally to the Brain age compared to DWMH (0.083 *vs* 0.002).

## Discussion

Establishment of age associated changes in magnitude, sequence and optimal number of MRI-determined neuroanatomic structure and white matter pathologies is imperative to develop a noninvasive quantitative precise clinical index for brain health and cognitive status. Here, we have investigated the temporal and spatial orders of neuroanatomic volumetry, small vessel pathology (white matter hyperintensity), and its kinetics with aging across subjects classified as cognitively normal (CN), cognitively impaired (CI), and cognitively impaired with an etiological diagnosis of AD (CI-AD). The brain structural changes in CN, CI, and CI-AD subjects were compared for the three age groups to identify early, intermediate, and late brain MRI events in the trajectory of aging. Using the brain MRI quantified features, a machine learning model is established to pinpoint an optimal minimum number of brain MRI segmented features discriminative of cognitive status. Moreover, for the first time a comprehensive architecture is developed for estimating the Brain Age using the neuroanatomic structures together with periventricular and deep white matter hyperintensity volume in aging brain.

The relatively slower GM and WM atrophy rate in the CI and CI-AD compared to the CN is because of EARLY loss of GM and WM volume in the CI and CI-AD subjects at the baseline age, as indicated from the lower intercept estimates and significantly lower volume from the age stratified volumetry analyses. The magnitude of GM and WM loss in CI and CI-AD subjects at early age is so remarkable that it attains plateau with aging. Quantitative estimation of gray and white matter at early age provides significant sensitivity for delineation of CI and CI-AD subjects from that of cognitively normal, but fails to distinguish between CI and CI-AD subjects. CI and CI-AD subjects have significantly overlapping GM and WM atrophy patterns. Although there was significant reduction of WM observed in CI and CI-AD subjects at early age groups, the differences were not observed at intermediate and late age groups plausibly because of faster WM loss in CN subjects compared to that of CI and CI-AD, owing to substantial loss at the early age groups of 50-64 years.

It is contradictory to notice reduced slope of structural atrophy in CI and CI-AD subjects in our study in contrast to prior reports of increased rate of loss of gray and white matter in AD subjects compared to the healthy control^35–37^. Given the substantial loss of gray and white matter being observed at much earlier age in the CI and CI-AD subjects in the current study, the prior reports of faster decline in GM and WM volume in cognitively impaired subjects is illusive. The rate of atrophy seems to depend on the baseline value, therefore for precise clinical sensitivity for assessing the brain health at a given age, the absolute volumetric quantification is imperative and its evaluation relative to the age matched normative library. We are developing an age matched normative reference of the majority of MRI determined brain structure volume.

Reduced hippocampal volume in CI and CI-AD subjects is not distinctive of CI and CI-AD at early age groups, similar to the pattern observed for the gray matter volume. The early onset AD cases (EOAD) i.e., before age of 65 lack involvement of memory components^38^, which is consistent with the findings of no significant difference in hippocampus, gray matter and white matter volume between CI and CI-AD subjects at the early age groups. The substantial loss in hippocampus provides sensitivity to differentiate between CI and CI-AD at the intermediate age group of ≥ 65 years at which almost 94% of late onset of AD cases are observed and become symptomatic.

The entorhinal cortical (EC) thickness marks a feature distinctive of CI and CI-AD subjects at the intermediate age group but lacks sensitivity to distinguish CN *vs* CI and CI *vs* CI-AD at the early age groups suggestive of entorhinal cortical thinning as a feature of LOAD than EOAD. We clearly observe a significant thinning of EC in CI and CI-AD cases compared to CN at intermediate and late age groups. However, a prior study that measured thickness of the entorhinal cortex did not observe significant difference between cognitively normal and amnestic MCI^39^. Thinning of entorhinal cortices are age dependent, therefore it is likely that the prior study did not observe thinning of EC because the measurements of EC thickness were pooled together irrespective of age. Interestingly, thinning of parahippocampal gyrus provides one of the unique early changes discriminative of CI *vs* CI-AD. Indeed, the medial cortical lobes are the primary site for tau deposition, a key characteristic in the manifestation of Alzheimer’s disease. Indeed, the prior measurements conducted to estimate the cortical thinning suggests parahippocampal cortical thinning, a remarkable feature of amnestic MCI and AD^39,40^.

Substantial hypertrophy of lateral ventricles serves as a unique early discriminator of CI-AD *vs* CI, CI *vs* CN, CI-AD *vs* CN subjects (Supplementary Table S3, S5). Prior studies indeed have discussed ventricular volume as a measure of disease progression in AD and MCI^41,42^. Although the CSF increase in CI-AD subjects is distinct from CI at the early age group, the CSF increase lacks sensitivity to distinguish CN vs CI. This indicates that a combination of CSF and ventricular volume must be considered together to monitor the aging associated cognitive disorders.

Progressive increase in WMH load with age reveals microvascular pruning with aging as an unavoidable vascular insult. Indeed, the vascular insult is increased in cognitively impaired groups compared to the CN as elevated WMH load in the CI and CI-AD groups is remarkably higher across intermediate and late age groups. Apparently, WMH load is not distinctive of CI and CI-AD subjects suggesting a common pathogenic underpinning of vascular components for MCI and AD subjects. Indeed, the faster rate of increase in WMH load in CI and CI-AD subjects compared to the CN indicates that WMH load beyond a threshold of normal aging changes may contribute towards the pathogenesis of aging-associated brain disorders. Loss of gray and white matter volume in CI and CI-AD subjects compared to the CN may elicit the punctate WMH load towards confluence and abrupt WMH increase, as observed from the exponential increase of WMH with aging. Gray matter (Neuronal body) and white matter (fiber) loss is an early event of compromised structural health, and beyond a threshold of WMH load when it tips up suddenly, the GM and WM loss may get accelerated resulting in accelerated cognitive decline.

Performing comprehensive whole brain quantitative regional volumetry is often not pragmatic in a clinical setting. Volumetry comparisons at the early, intermediate and late age groups (Supplementary Table S3, S5) revealed that not all brain structural and microvascular changes serve as a discriminator for cognitive status, as the magnitude of changes across events is subtle, and follow a unique sequence which are temporally and spatially separated. Therefore, it is impeccable to deduce the minimum number of optimal, unique, easy-to-quantify MRI features distinctive of cognitive health. Three distinct features comprising a structural feature i.e., the total brain volume (= gray matter and white matter), a glymphatic system as CSF volume and a microvascular feature i.e., white matter hyperintensity (WMH) provided a significantly higher accuracy for predicting the cognitive status. Inclusion of additional MRI determined neuroanatomical structures did not improve the accuracy, and led to plateauing of the accuracy.

Inclusion of quantitative WMH lesion load in neurobiological understanding of brain and cognitive health during aging and aging associated pathologies is poorly studied. Given WMH presence in almost all the brains studied in this study across CN, CI and CI-AD, in miniscule or high amounts, it is imperative to deduce its implications in determining brain health. The ML model clearly indicates that WMH volume is one of the unique features along with total brain volume and CSF, as a predictor of clinical cognitive status. WMH quantitation along with structural volumetry will provide the extent of vascular insult and its impact on structure and functional status of the brain. Our brain age model is the first report wherein load of periventricular and deep white matter hyperintensity in subjects with low and high white matter hyperintensity revealed a clearly distinct higher brain age gap. Subjects with WMH >5 ml had brain age gaps more than ∼3 years compared to the subjects with low or miniscule WMH. This further establishes that in order to understand the normal aging and pathological aging trajectory, one cannot treat the subjects with WMH and without WMH together. Owing to the vascular insular from WMH, the structural atrophy and hypertrophy kinetics will be distinct depending upon WMH load.

Brain age is indeed a comprehensive representation of a set of neuroanatomical quantities which are major contributors, such as 3rd Ventricle volume estimates with chronological aging contributes most towards brain age with an importance factor of 0.14 out of the top 10 brain features such as volume of accumbens, brain segmented volume by total intracranial volume ratio, amygdala volume, post-central thickness, cerebellum white matter volume, caudal-anterior cingulate white matter volume, posterior cingulate surface area, temporal pole white matter volume, and post-central white matter volume (Fig.S4 A). It is interesting to further enhance the biological underpinning of these top 10 features which appears to be key players for brain age and brain health in the subjects with no WMH deposition. It is intriguing to note that a unique set of brain features contribute towards the brain age for the subjects with high WMH load compared to the low/nil WMH subjects (Fig.S4 B). Despite the differences in the set of important features between the two groups, a few features were common contributors of the Brain Age estimation. Therefore, here we establish that brain age gap may serve as a potential clinical measure of brain health, when investigated into structural and commonly observed small vessel pathology of WMH.

## CONCLUSION

‘Three’ unique brain changes associated with aging, wherein two structural features Total brain volume and CSF volume, together with WMH lesion load provides highly precise discrimination of cognitive status as cognitively normal (CN), impaired (CI) and AD (CI-AD). The unique brain age model that uses WMH load along with neuroanatomic quantities depicts that the elevated load of WMH at a given chronological age contributes to an increased Brain Age gap even in subjects screened as cognitively normal. Despite the presence of a structural lesion in the form of WMH in the brain, an individual gets classified as cognitively normal, calling for a debate on the precision of the clinical cognitive evaluations as a true indicator of brain health. Increased Brain Age gap with elevated WMH load even at the early age groups is suggestive of profound vascular insult resulting in early neuroanatomic atrophy or hypertrophy.

Moreover, upon comprehensive quantification of neuroanatomic and WMH volume, this study establishes that a unique sequence and magnitude of structural and microvascular features have early, intermediate and late age sensitivity to delineate the cognitive status. Wherein, the ventricular volume serves as an early feature distinctive between all the three cognitive groups while loss of hippocampus volume, gray matter and white matter volume is distinctive of CI and CI-AD from CN in the early age group but lacks sensitivity to discriminate between CI and CI-AD. It is remarkable to note that medial cortical thinning from Parahippocampal gyrus is an Early event discriminative between cognitively impaired subjects because of MCI (CI) from that of cognitive impairment due to Alzheimer’s disease (CI-AD). Higher load of WMH and faster kinetics in CI and CI-AD subjects indicates that WMH deposition may be one of the contributors that would affect the chronological aging pattern of structural atrophy and hypertrophy. Indeed, WMH serves as one of the three unique features discriminative of cognitive status, and also estimates of PVWMH load is a significant contributor towards increased brain age gap. The overall findings from the study advocates that quantification and analyses of a unique set of brain structural volume together with microvascular pathology of WMH may serve as a non-invasive potential clinically precise signature for further biological underpinning of aging and cognitive impairment.

## Data availability

Data used in the current study are available through the NACC (naccdata.org) and ADNI website (https://adni.loni.usc.edu). Detailed data analysis plan and syntax will be made available upon request to the corresponding author.

## Acknowledgments

The Brain age estimation model was established using neuroanatomic data from the National Alzheimer’s Coordinating Centre (NACC). The Brain age modeling and the estimation of Brain age gap was cross-validated using the data from Alzheimer’s Disease Neuroimaging Initiative (ADNI). The NACC database is funded by NIA/NIH Grant U24 AG072122. NACC data are contributed by the NIA-funded ADRCs: P30 AG062429 (PI James Brewer, MD, PhD), P30 AG066468 (PI Oscar Lopez, MD), P30 AG062421 (PI Bradley Hyman, MD, PhD), P30 AG066509 (PI Thomas Grabowski, MD), P30 AG066514 (PI Mary Sano, PhD), P30 AG066530 (PI Helena Chui, MD), P30 AG066507 (PI Marilyn Albert, PhD), P30 AG066444 (PI John Morris, MD), P30 AG066518 (PI Jeffrey Kaye, MD), P30 AG066512 (PI Thomas Wisniewski, MD), P30 AG066462 (PI Scott Small, MD), P30 AG072979 (PI David Wolk, MD), P30 AG072972 (PI Charles DeCarli, MD), P30 AG072976 (PI Andrew Saykin, PsyD), P30 AG072975 (PI David Bennett, MD), P30 AG072978 (PI Neil Kowall, MD), P30 AG072977 (PI Robert Vassar, PhD), P30 AG066519 (PI Frank LaFerla, PhD), P30 AG062677 (PI Ronald Petersen, MD, PhD), P30 AG079280 (PI Eric Reiman, MD), P30 AG062422 (PI Gil Rabinovici, MD), P30 AG066511 (PI Allan Levey, MD, PhD), P30 AG072946 (PI Linda Van Eldik, PhD), P30 AG062715 (PI Sanjay Asthana, MD, FRCP), P30 AG072973 (PI Russell Swerdlow, MD), P30 AG066506 (PI Todd Golde, MD, PhD), P30 AG066508 (PI Stephen Strittmatter, MD, PhD), P30 AG066515 (PI Victor Henderson, MD, MS), P30 AG072947 (PI Suzanne Craft, PhD), P30 AG072931 (PI Henry Paulson, MD, PhD), P30 AG066546 (PI Sudha Seshadri, MD), P20 AG068024 (PI Erik Roberson, MD, PhD), P20 AG068053 (PI Justin Miller, PhD), P20 AG068077 (PI Gary Rosenberg, MD), P20 AG068082 (PI Angela Jefferson, PhD), P30 AG072958 (PI Heather Whitson, MD), P30 AG072959 (PI James Leverenz, MD).Data collection and sharing for the ADNI data was funded by the Alzheimer’s Disease Neuroimaging Initiative (ADNI) (National Institutes of Health Grant U01 AG024904) and DOD ADNI (Department of Defense award number W81XWH-12-2-0012).

ADNI is funded by the National Institute on Aging, the National Institute of Biomedical Imaging and Bioengineering, and through generous contributions from the following: AbbVie, Alzheimer’s Association; Alzheimer’s Drug Discovery Foundation; Araclon Biotech; BioClinica, Inc.; Biogen; Bristol-Myers Squibb Company; CereSpir, Inc.; Cogstate; Eisai Inc.; Elan Pharmaceuticals, Inc.; Eli Lilly and Company; EuroImmun; F. Hoffmann-La Roche Ltd and its affiliated company Genentech, Inc.; Fujirebio; GE Healthcare; IXICO Ltd.; Janssen Alzheimer Immunotherapy Research & Development, LLC.; Johnson & Johnson Pharmaceutical Research & Development LLC.; Lumosity; Lundbeck; Merck & Co., Inc.; Meso Scale Diagnostics, LLC.; NeuroRx Research; Neurotrack Technologies; Novartis Pharmaceuticals Corporation; Pfizer Inc.; Piramal Imaging; Servier; Takeda Pharmaceutical Company; and Transition Therapeutics. The Canadian Institutes of Health Research is providing funds to support ADNI clinical sites in Canada. Private sector contributions are facilitated by the Foundation for the National Institutes of Health (www.fnih.org). The grantee organization is the Northern California Institute for Research and Education, and the study is coordinated by the Alzheimer’s Therapeutic Research Institute at the University of Southern California. ADNI data are disseminated by the Laboratory for NeuroImaging at the University of Southern California. We also acknowledge the contribution of Mr. V. Ashwin and Mr. Arkaprava Majumdar for helping in machine learning methods.

## Author information

### Authors and Affiliations

**Indian Institute of Science Education and Research (IISER) Berhampur, India.**

Neha Yadav, Niraj Kumar Gupta, Darshit Thakar & Vivek Tiwari

### Contributions

V.T. and N.Y. conceptualized and designed the study; N.Y., N.K.G., D.T. and V.T. analyzed the data, contributed to data interpretation; N.Y. and V.T. drafted the manuscript. All authors reviewed the results and approved the final version of the manuscript.

### Corresponding Author

Correspondence to Vivek Tiwari

## Ethics declarations

Conflict of interests

The authors declare no conflict of interests.

## Supplementary Figure

**Figure S1:**
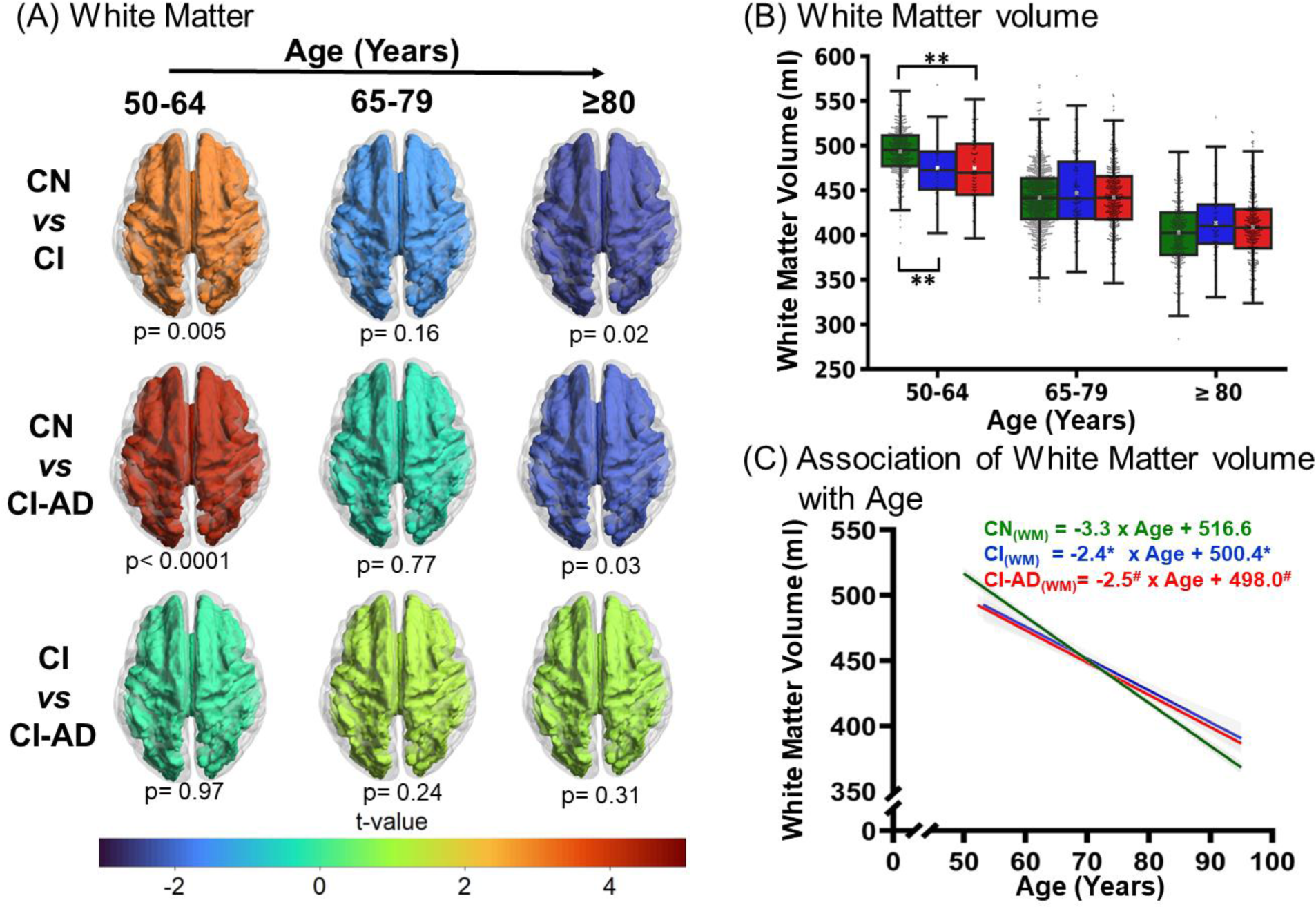
Magnitude and kinetics of White matter volume with age across CN, CI, and CI-AD subjects. **(A)** White matter volume comparison between CN *vs* CI, CN *vs* CI-AD, and CI *vs* CI-AD across three age groups i.e., 50-64 (early), 65-70 (intermediate), and ≥80 shown by the t-map where the color represents the t-value indicative of the difference between the two cognitive groups. Early loss of the white matter volume is observed for CI and CI-AD subjects compared to the CN subjects while CI-AD and CI subjects are not distinguishable (shown by the warmer color). The significance was set to p <0.016 (Bonferroni corrected). **(B)** The box plot depicts median (solid line) and mean (white square) volume of white matter in CN, CI, and CI-AD subjects across early, intermediate, and late age groups. White matter volume between the cognitive groups was compared using unpaired, two-tailed Welch’s *t*-test followed by Bonferroni correction. Statistical significance is depicted as * p<0.016, **p<0.001. **(C)** Linear regression of volume of white matter (WM) with age. The linear regression was performed upon setting up the intercept at 50 years for CN (green), CI (blue), and CI-AD (red) subjects. Statistical significance for the slope and intercept comparison between CN vs CI (*), CN vs CI-AD (^#^), and CI vs CI-AD (^$^) was set at p < 0.05.

**Figure S2:**
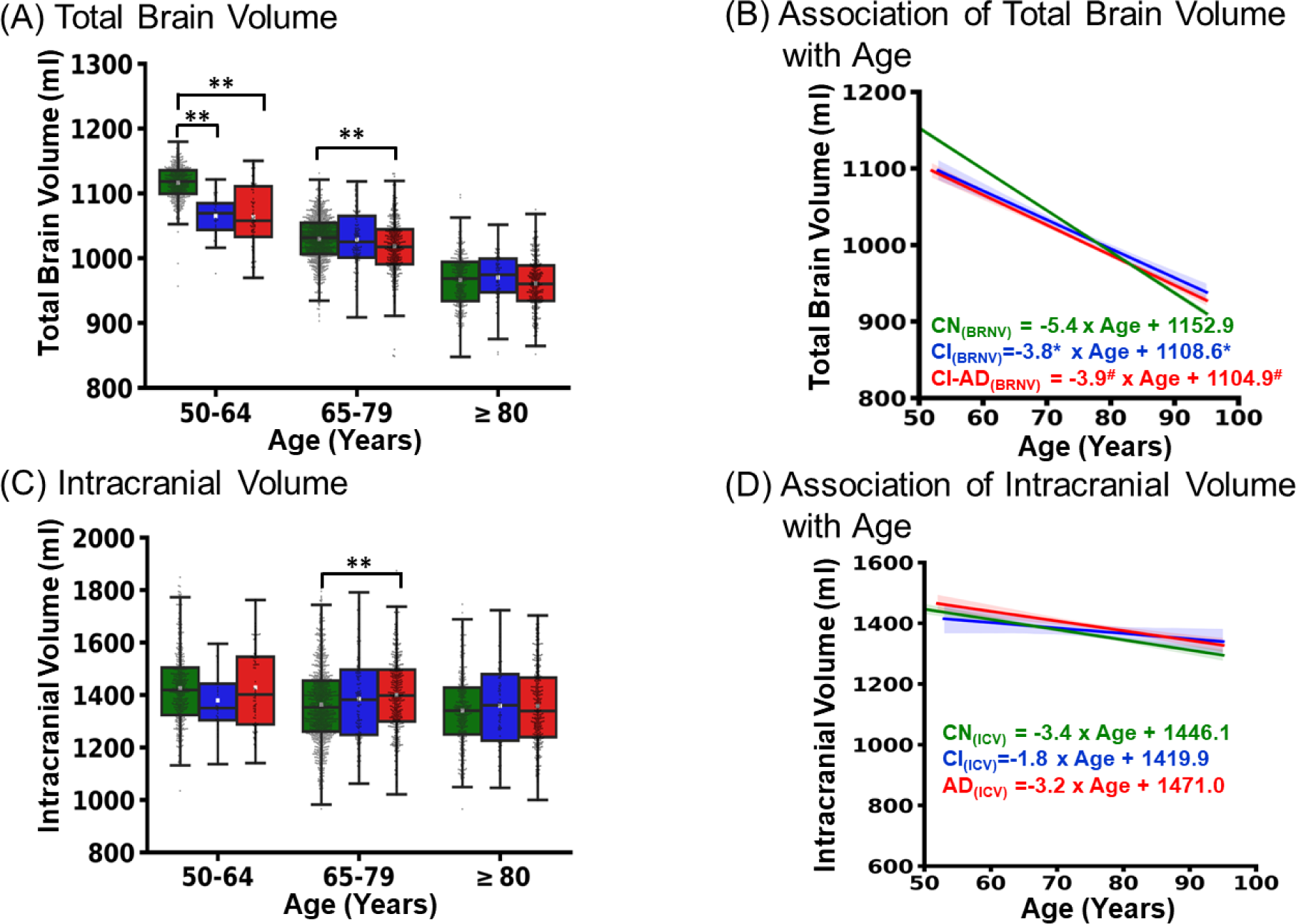
**(A)** Boxplot illustrating the total brain volume across early, intermediate and late age groups for CN (green), CI (blue), and CI-AD (red) subjects. **(B)** Linear regression of Total Brain volume with age upon setting up the intercept at 50 years. **(C)** Boxplot depicts Total Intracranial volume across early), intermediate, and late age groups for CN (green), CI (blue), and CI-AD (red) subjects**. (D)** Linear regression of Total Intracranial volume with age upon setting up the intercept at 50 years. Statistical significance for the slope and intercept comparison between CN vs CI (*), CN vs CI-AD (^#^), and CI vs CI-AD (^$^) was set at p <0.05. The mean volumes were compared using unpaired two-tailed t-test followed by Bonferroni correction. Statistical significance for the mean volume comparison among cognitive groups (CN, CI, and CI-AD) across the stratified age groups is depicted as * p <0.016, **p <0.001

**Figure S3:**
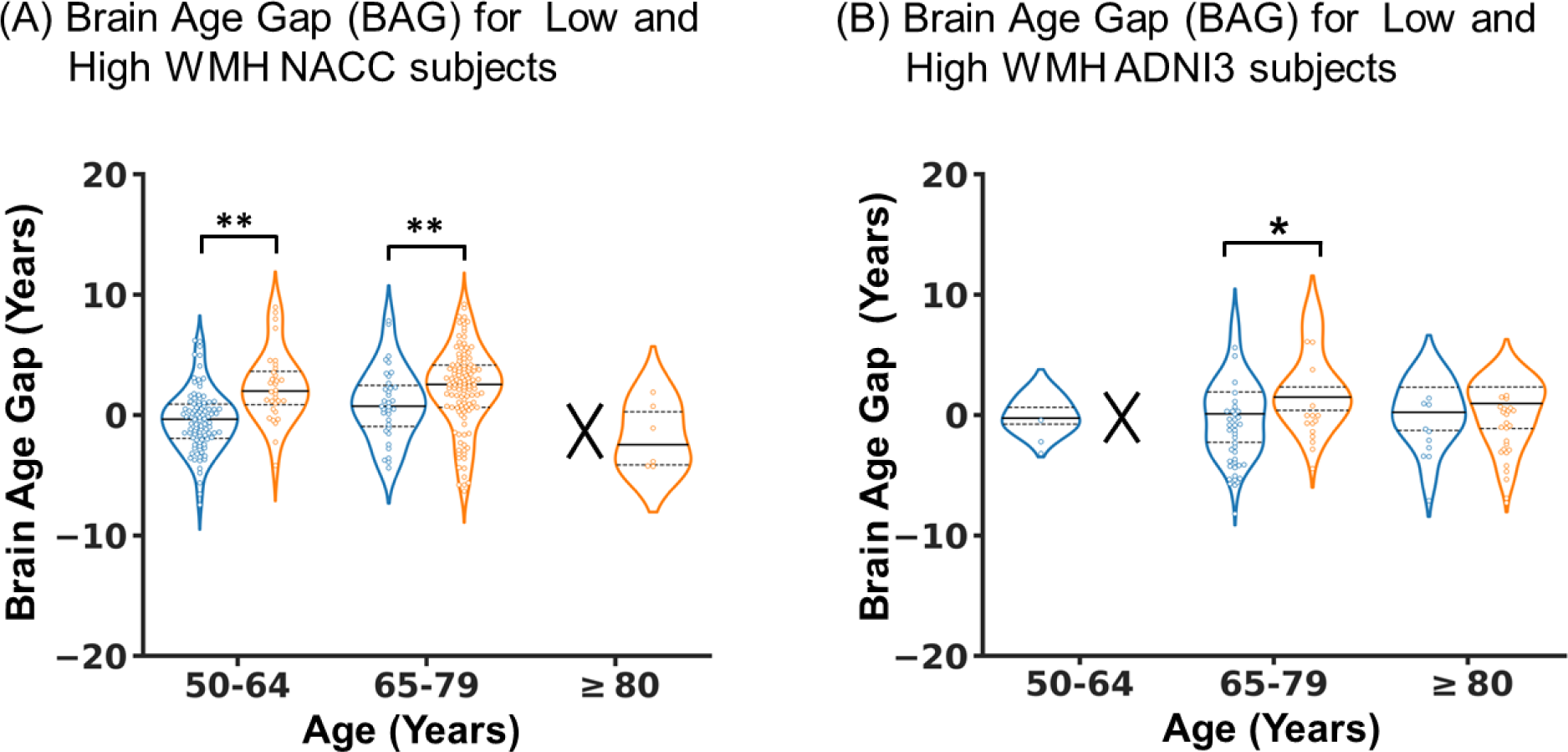
**(A)** Brain Age Gap (BAG) of the cognitively normal NACC subjects in early age group (50-64) with no or low WMH <1.5 ml (blue, n=95) and high WMH 5-10 ml (orange, n=32), intermediate age group (65-79) with no or low WMH (blue, n=35) and high WMH (orange, n=118), and late age group (≥80) with no or low WMH (blue, n=0) and high WMH (orange, n=13). **(B)** Brain Age Gap (BAG) for the cognitively normal subjects from ADNI3 cohort in early age group with no or low WMH (blue, n=3) and high WMH (orange, n=0), intermediate age group with no or low WMH (blue, n=39) and high WMH (orange, n=15), and late age group with no or low WMH (blue, n=10) and high WMH (orange, n=25). Continuous line, median; dotted line, interquartile range (IQR). Statistical significance by unpaired two-tailed *t*-test for the comparison between group having no or low WMH and high WMH in the early, intermediate, and late age groups is depicted as * p <0.05, **p <0.001.

**Figure S4:**
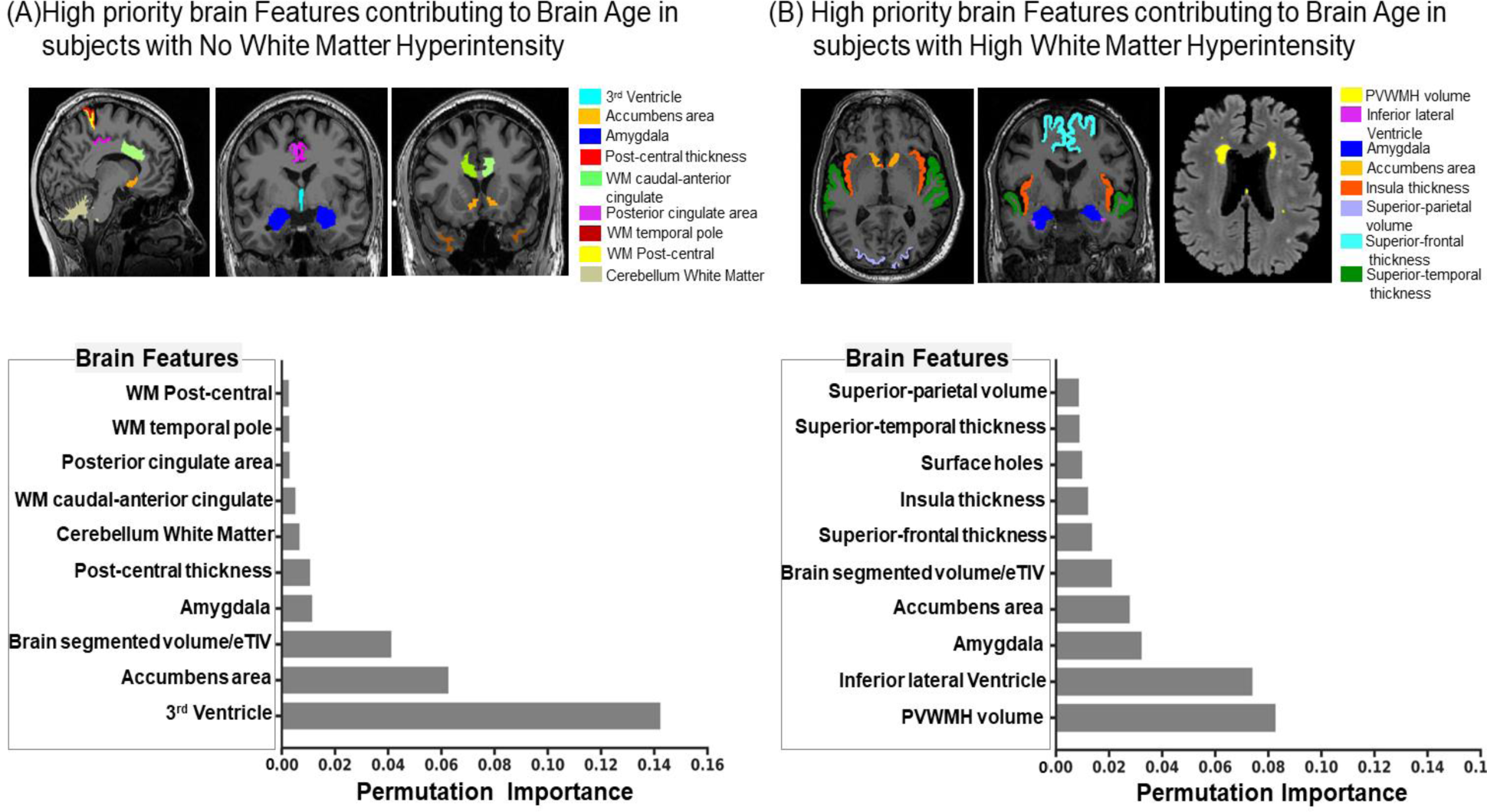
The relative contribution of Neuroanatomic Structures towards Brain Age Estimation. The top ten brain MRI determined neuroanatomic quantities ranked using permutation importance in decreasing order of contribution towards Brain Age estimation for subjects with (A) no WMH (PVWMH and DWMH <1.5 ml) and subjects with (B) High WMH (PVWMH + DWMH > 5 ml). The Coronal, sagittal and axial MRI slices depicting segmented masks of important brain features overlaid on T1-weighted and T2 -FLAIR images. The permutation importance reflects quantitative reduction in the weight of the brain age estimation model performance when performed in absence of the respective feature. eTIV: estimated Total Intracranial Volume.

## Supplementary Tables

**Supplementary Table S1.** Demographics of the cohort.

| Cognitive Status | Number of participants (M/F) | Age participants (mean ± SD) | Education (mean ± SD) | Number of MRI scans (M/F) | Age MRI Scans (mean ± SD) |
| --- | --- | --- | --- | --- | --- |
| CN | 1082<br>(367/715) | 70.8 ± 10.0 | 15.8 ± 5.4 | 1616<br>(537/1079) | 71.3 ± 9.7 |
| CI | 197<br>(93/104) | 74.3 ± 9.2 | 15.1 ± 7.2 | 252<br>(127/125) | 74.8 ± 9.0 |
| CI-AD | 588<br>(312/276) | 76.1 ± 8.2 | 15.5 ± 7.0 | 744<br>(404/370) | 76.6 ± 8.6 |

SD- Standard deviation
M- Male
F- Female

**Supplementary Table S2 (A).** Global Hypothesis using nonparametric test with the three cognitive status as factor and eight neuroanatomic structures as response variables.

| Age Group | Nonparametric test | Test Statistic | df1 | df2 | p-value | Permutation Test p-value |
| --- | --- | --- | --- | --- | --- | --- |
| **50 - 64** | ANOVA type test p-value | 22.31 | 8.09 | 785.49 | 0 | 0 |
|  | McKeon approx. for the Lawley Hotelling Test | 83.19 | 16.00 | 839.15 | 0 | 0 |
|  | Muller approx. for the Bartlett-Nanda-Pillai Test | 63.34 | 16.09 | 1035.92 | 0 | 0 |
|  | Wilks Lambda | 72.96 | 16.00 | 1030.00 | 0 | 0 |
| **65 -79** | ANOVA type test p-value | 41.39 | 7.03 | 2866.74 | 0 | 0 |
|  | McKeon approx. for the Lawley Hotelling Test | 213.54 | 16.00 | 2257.87 | 0 | 0 |
|  | Muller approx. for the Bartlett-Nanda-Pillai Test | 100.17 | 16.03 | 2769.97 | 0 | 0 |
|  | Wilks Lambda | 151.99 | 16.00 | 2764.00 | 0 | 0 |
| **≥80** | ANOVA type test p-value | 16.33 | 7.98 | 1915.59 | 0 | 0 |
|  | McKeon approx. for the Lawley Hotelling Test | 27.76 | 16.00 | 1166.42 | 0 | 0 |
|  | Muller approx. for the Bartlett-Nanda-Pillai Test | 21.95 | 16.07 | 1435.94 | 0 | 0 |
|  | Wilks Lambda | 24.83 | 24.83 | 1430.00 | 0 | 0 |

**Supplementary Table S2 (B).** Nonparametric relative effect of cognitive status on brain regions.

| Age Group | Cognitive Status | Gray Matter Volume | White Matter Volume | CSF | Hippocampus volume | Lateral Ventricle Volume | Para-hippocampal Gyrus Thickness | Entorhinal Cortex Thickness | Total Brain Volume |
| --- | --- | --- | --- | --- | --- | --- | --- | --- | --- |
| 50 - 64 | CN | 0.57 | 0.53 | 0.45 | 0.55 | 0.44 | 0.55 | 0.54 | 0.57 |
|  | CI | 0.22 | 0.36 | 0.47 | 0.36 | 0.60 | 0.49 | 0.46 | 0.18 |
|  | CI-AD | 0.25 | 0.38 | 0.76 | 0.27 | 0.76 | 0.26 | 0.31 | 0.26 |
| 65 -79 | CN | 0.54 | 0.50 | 0.40 | 0.59 | 0.41 | 0.57 | 0.59 | 0.52 |
|  | CI | 0.49 | 0.50 | 0.53 | 0.50 | 0.58 | 0.47 | 0.51 | 0.51 |
|  | CI-AD | 0.42 | 0.52 | 0.70 | 0.31 | 0.67 | 0.34 | 0.29 | 0.44 |
| ≥ 80 | CN | 0.56 | 0.47 | 0.44 | 0.61 | 0.41 | 0.59 | 0.63 | 0.51 |
|  | CI | 0.50 | 0.55 | 0.48 | 0.52 | 0.47 | 0.52 | 0.52 | 0.55 |
|  | CI-AD | 0.43 | 0.52 | 0.57 | 0.37 | 0.61 | 0.40 | 0.35 | 0.47 |

**Supplementary Table S3.**
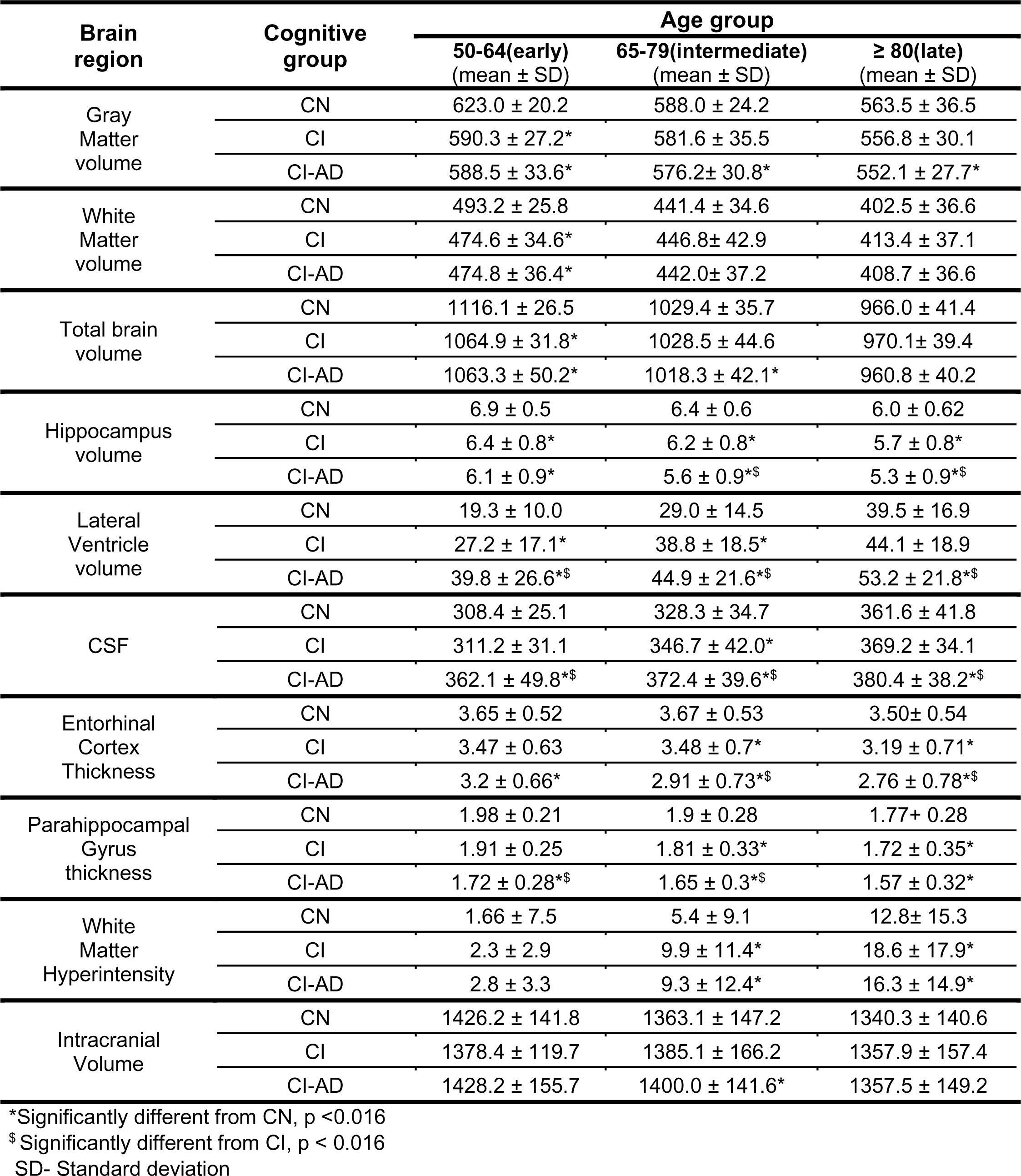
Age groupwise mean volume/thickness of brain regions.

| Brain region | Cognitive group | Age group |  |  |
| --- | --- | --- | --- | --- |
| | | 50-64(early)<br>(mean ± SD) | 65-79(intermediate)<br>(mean ± SD) | $\geq 80$(late)<br>(mean ± SD) |
| Gray Matter volume | CN | 623.0 ± 20.2 | 588.0 ± 24.2 | 563.5 ± 36.5 |
| | CI | 590.3 ± 27.2* | 581.6 ± 35.5 | 556.8 ± 30.1 |
| | CI-AD | 588.5 ± 33.6* | 576.2 ± 30.8* | 552.1 ± 27.7* |
| White Matter volume | CN | 493.2 ± 25.8 | 441.4 ± 34.6 | 402.5 ± 36.6 |
| | CI | 474.6 ± 34.6* | 446.8 ± 42.9 | 413.4 ± 37.1 |
| | CI-AD | 474.8 ± 36.4* | 442.0 ± 37.2 | 408.7 ± 36.6 |
| Total brain volume | CN | 1116.1 ± 26.5 | 1029.4 ± 35.7 | 966.0 ± 41.4 |
| | CI | 1064.9 ± 31.8* | 1028.5 ± 44.6 | 970.1 ± 39.4 |
| | CI-AD | 1063.3 ± 50.2* | 1018.3 ± 42.1* | 960.8 ± 40.2 |
| Hippocampus volume | CN | 6.9 ± 0.5 | 6.4 ± 0.6 | 6.0 ± 0.62 |
| | CI | 6.4 ± 0.8* | 6.2 ± 0.8* | 5.7 ± 0.8* |
| | CI-AD | 6.1 ± 0.9* | 5.6 ± 0.9*\$ | 5.3 ± 0.9*\$ |
| Lateral Ventricle volume | CN | 19.3 ± 10.0 | 29.0 ± 14.5 | 39.5 ± 16.9 |
| | CI | 27.2 ± 17.1* | 38.8 ± 18.5* | 44.1 ± 18.9 |
| | CI-AD | 39.8 ± 26.6*\$ | 44.9 ± 21.6*\$ | 53.2 ± 21.8*\$ |
| CSF | CN | 308.4 ± 25.1 | 328.3 ± 34.7 | 361.6 ± 41.8 |
| | CI | 311.2 ± 31.1 | 346.7 ± 42.0* | 369.2 ± 34.1 |
| | CI-AD | 362.1 ± 49.8*\$ | 372.4 ± 39.6*\$ | 380.4 ± 38.2*\$ |
| Entorhinal Cortex Thickness | CN | 3.65 ± 0.52 | 3.67 ± 0.53 | 3.50 ± 0.54 |
| | CI | 3.47 ± 0.63 | 3.48 ± 0.7* | 3.19 ± 0.71* |
| | CI-AD | 3.2 ± 0.66* | 2.91 ± 0.73*\$ | 2.76 ± 0.78*\$ |
| Parahippocampal Gyrus thickness | CN | 1.98 ± 0.21 | 1.9 ± 0.28 | 1.77 ± 0.28 |
| | CI | 1.91 ± 0.25 | 1.81 ± 0.33* | 1.72 ± 0.35* |
| | CI-AD | 1.72 ± 0.28*\$ | 1.65 ± 0.3*\$ | 1.57 ± 0.32* |
| White Matter Hyperintensity | CN | 1.66 ± 7.5 | 5.4 ± 9.1 | 12.8 ± 15.3 |
| | CI | 2.3 ± 2.9 | 9.9 ± 11.4* | 18.6 ± 17.9* |
| | CI-AD | 2.8 ± 3.3 | 9.3 ± 12.4* | 16.3 ± 14.9* |
| Intracranial Volume | CN | 1426.2 ± 141.8 | 1363.1 ± 147.2 | 1340.3 ± 140.6 |
| | CI | 1378.4 ± 119.7 | 1385.1 ± 166.2 | 1357.9 ± 157.4 |
| | CI-AD | 1428.2 ± 155.7 | 1400.0 ± 141.6* | 1357.5 ± 149.2 |

\*Significantly different from CN, p < 0.016
\$ Significantly different from CI, p < 0.016
SD- Standard deviation

**Supplementary Table S4.** Multivariate linear regression model coefficients and standard error for brain region volume/thickness.

| Brain region | $\beta_0$ | | $\beta_1$ | | $\beta_2$ | | $\beta_3$ | | $\beta_4$ | | $\beta_5$ | |
| --- | --- | --- | --- | --- | --- | --- | --- | --- | --- | --- | --- | --- |
|  | Value | SE | Value | SE | Value | SE | Value | SE | Value | SE | Value | SE |
| Gray matter Volume | 636.2** | 1.6 | -2.1** | 0.07 | 0.7** | 0.2 | -28.0** | 5.2 | 0.6** | 0.13 | -29.3** | 3.5 |
| White Matter Volume | 516.6** | 2.1 | -3.3** | 0.09 | 0.9** | 0.25 | -16.2* | 6.7 | 0.8** | 0.17 | -18.6** | 4.5 |
| Total Brain volume | 1152.9** | 2.24 | -5.4** | 0.1 | 1.6** | 0.28 | -44.3** | 7.3 | 1.4** | 0.18 | -48.0** | 4.89 |
| Hippocampus volume | 7.14** | 0.04 | -0.032** | 0.0018 | 0.006 | 0.0053 | -0.41** | 0.139 | 0.007 | 0.003 | -0.95** | 0.093 |
| Lateral ventricle | 12.4** | 1.02 | 0.8** | 0.04 | -0.1 | 0.13 | 11.0** | 3.33 | -0.2** | 0.08 | 20.9** | 2.2 |
| CSF | 284.6** | 2.12 | 2.1** | 0.09 | 0.06 | 0.26 | 9.9 | 6.9 | -1.3** | 0.2 | 67.1** | 4.6 |
| Parahippocampal gyrus thickness | 2.06** | 0.016 | -0.008** | 0.0007 | 0.002 | 0.002 | -0.10 | 0.055 | 0.002 | 0.001 | -0.27** | 0.036 |
| Entorhinal cortex thickness | 3.73** | 0.04 | -0.005** | 0.001 | -0.01* | 0.004 | 0.02 | 0.12 | -0.01** | 0.003 | -0.46** | 0.08 |
| Intracranial Volume | 1446.1** | 8.7 | -3.4** | 0.4 | 1.6 | 1.1 | -26.2 | 28.5 | 0.2 | 0.71 | 24.9 | 19.1 |

\*p<0.05, \*\*p<0.01 $\beta_0$ - Intercept for Control group (CN), $\beta_1$ - coefficient of Age for CN, $\beta_2$ - coefficient for Age and CI interaction, $\beta_3$ – coefficient for CI, $\beta_4$ – coefficient for AD and Age interaction, $\beta_5$ – coefficient for AD. CI =1 for CI group otherwise 0, AD=1 for CI-AD group otherwise 0. SE- Standard error in the estimated values of the coefficient

**Supplementary Table S5.**
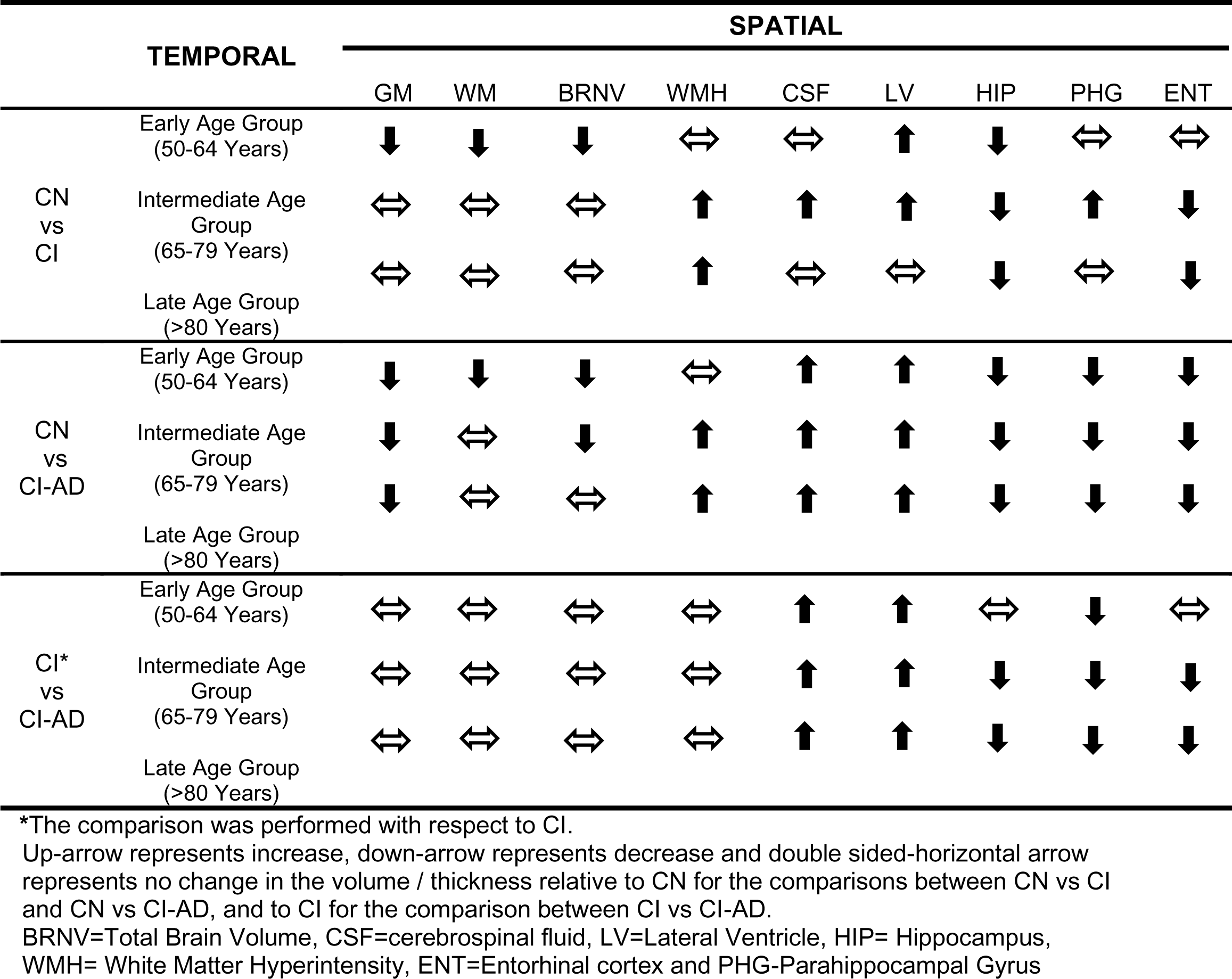
Temporal and spatial Order of Events in Aging and Cognitive Impairment.

| TEMPORAL |  | SPATIAL |  |  |  |  |  |  |  |  |
| --- | --- | --- | --- | --- | --- | --- | --- | --- | --- | --- |
|  |  | GM | WM | BRNV | WMH | CSF | LV | HIP | PHG | ENT |
| CN<br>vs<br>CI | Early Age Group<br>(50-64 Years) | ↓ | ↓ | ↓ | ↔ | ↔ | ↑ | ↓ | ↔ | ↔ |
|  | Intermediate Age<br>Group<br>(65-79 Years) | ↔ | ↔ | ↔ | ↑ | ↑ | ↑ | ↓ | ↑ | ↓ |
|  | Late Age Group<br>(>80 Years) | ↔ | ↔ | ↔ | ↑ | ↔ | ↔ | ↓ | ↔ | ↓ |
| CN<br>vs<br>CI-AD | Early Age Group<br>(50-64 Years) | ↓ | ↓ | ↓ | ↔ | ↑ | ↑ | ↓ | ↓ | ↓ |
|  | Intermediate Age<br>Group<br>(65-79 Years) | ↓ | ↔ | ↓ | ↑ | ↑ | ↑ | ↓ | ↓ | ↓ |
|  | Late Age Group<br>(>80 Years) | ↓ | ↔ | ↔ | ↑ | ↑ | ↑ | ↓ | ↓ | ↓ |
| CI*<br>vs<br>CI-AD | Early Age Group<br>(50-64 Years) | ↔ | ↔ | ↔ | ↔ | ↑ | ↑ | ↔ | ↓ | ↔ |
|  | Intermediate Age<br>Group<br>(65-79 Years) | ↔ | ↔ | ↔ | ↔ | ↑ | ↑ | ↓ | ↓ | ↓ |
|  | Late Age Group<br>(>80 Years) | ↔ | ↔ | ↔ | ↔ | ↑ | ↑ | ↓ | ↓ | ↓ |

\*The comparison was performed with respect to CI.
Up-arrow represents increase, down-arrow represents decrease and double sided-horizontal arrow represents no change in the volume / thickness relative to CN for the comparisons between CN vs CI and CN vs CI-AD, and to CI for the comparison between CI vs CI-AD.
BRNV=Total Brain Volume, CSF=cerebrospinal fluid, LV=Lateral Ventricle, HIP= Hippocampus, WMH= White Matter Hyperintensity, ENT=Entorhinal cortex and PHG-Parahippocampal Gyrus

